# Fixed-confidence identification of combination interventions in stochastic gene-network models, with a benchmark and an application to acute myeloid leukemia

**DOI:** 10.64898/2026.11.22.768439

**Authors:** Wasim Raza, Saba Mehmood, Carsten Enderlein, Fabrício Alves Barbosa da Silva

**Affiliations:** Developmental Biology and Dynamical Systems Group, Universidade Federal do Rio de Janeiro, Professor Geraldo Cidade Campus, Duque de Caxias, Rio de Janeiro, Brazil; Department of Mathematics, University of Management and Technology (UMT), Lahore, Pakistan; NUMPEX-Nano, Campus Duque de Caxias da UFRJ, Rio de Janeiro, Brazil; Scientific Computing Programme, Oswaldo Cruz Foundation (FIOCRUZ), Rio de Janeiro, Brazil; Department of Mathematics, Faculty of Sciences, Thal University, Bhakkar, Pakistan

**Author notes:** Corresponding author: Fabrício Alves Barbosa da Silva.

## Abstract

Mechanistic gene-network models are widely used to rank drug-target combinations. Each ranking rests on a finite number of noisy simulations, yet the confidence that the top-ranked set is truly best in the model is rarely stated. We formulate intervention search in a stochastic network model as fixed-confidence best-arm identification and give certified search: empirical-Bernstein confidence sequences that return an *ε*-optimal intervention set with probability at least 1 − 2*δ* under any sampling rule, with successive elimination and batch-parallel LUCB as sampling rules. We compare them with uniform sampling, successive halving, greedy forward selection, sequential LUCB and UGapE, and ablate each design choice, on four networks (a BeatAML-calibrated AML network and three published cancer Boolean models; 1,275 to 5,488 candidate sets) under a protocol written before the runs. All 21 certified benchmark runs returned an *ε*-optimal set (exact one-sided 95% lower bound 86.7%), as did all 4,560 and 1,980 replayed runs of the comparisons, using 6.4 to 19.0 times fewer simulations than uniform sampling with the same guarantee. Elimination and the variance-adaptive radius account for most of the saving (2.1 to 34.1 and 1.3 to 3.9 times). Sequential LUCB and UGapE use fewer simulations in total but are slow on parallel hardware; batch-parallel LUCB, with as many candidates per round as workers, was at most 1% slower than successive elimination in any setting and up to 3.2 times faster, and on the three published networks used 2.0 to 3.1 times fewer simulations. Applied to AML, a search over eight drug classes returns ibrutinib + midostaurin + an ERK-inhibitor class (model score 0.710 against 0.448 untreated), which is the best set in all 15 related calibrations of the network and exceeds additive and Bliss-independence expectations.

## 1 Introduction

Mechanistic gene-network models are widely used to suggest drug targets. A drug is represented by forcing its target genes off, the network is simulated, and the intervention that pushes the system furthest from a malignant state toward an apoptotic one is ranked first [1, 2, 3, 4]. Two features of this workflow matter for how far a ranking can be trusted. Many of the models are stochastic, either by construction (probabilistic Boolean networks [5]) or through added noise, so the score of an intervention is an expectation that can only be estimated. And the number of candidates is large: with *N* targetable genes there are 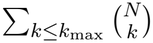 sets to compare, so naming a winner means comparing thousands of noisy estimates. The pipeline we build on [4] draws about thirty trajectories per candidate and reports the empirical maximum, without a statement of how likely that maximum is to be the true best.

Attractor-based control of Boolean networks is well developed [6] and addresses a different question, namely how to drive a network to a chosen attractor. The question here is how many simulations are needed before one can state, with a given confidence, that a candidate intervention is within *ε* of the best in a finite list.

This is fixed-confidence best-arm identification, for which successive elimination, LUCB and UGapE offer finite-sample guarantees [7, 8, 9, 10]. Applying it to intervention search raises three practical questions that we address. First, which design choices make a certificate affordable for bounded simulation scores: elimination, the confidence radius, the sampling schedule? Second, how does a certified search compare with the simpler procedures used in practice and with established best-arm algorithms, when all use the same confidence sequence? Third, how does its cost depend on the accuracy target and on the structure of the candidate family?

We make four contributions.

1. **Method**. Certified search is a certificate construction: empirical-Bernstein confidence sequences whose union bound over (candidate, number of batches) holds under any sampling rule, so an *ε*-optimal intervention set is returned with probability at least 1 *−* 2*δ* and runs that reach their sample cap are flagged as uncertified. We use two sampling rules, successive elimination (Algorithm 1; the rule of the pre-specified benchmark) and batch-parallel LUCB [11], and give an explicit-constant sample-complexity bound for successive elimination that we check numerically (Theorems 1–3).
2. **Comparison and ablation**. On a benchmark whose networks, objectives and success criteria were fixed before the runs, we compare certified search with uniform sampling, successive halving, greedy forward selection, LUCB and UGapE, in total simulations and in elapsed time on parallel workers, and ablate elimination, the radius, the checkpoint schedule and the batch size. The comparison is replayed on all four networks (13 settings), and a controlled experiment separates the effect of family size from that of near-ties.
3. **Audits of the guarantee**. Across 4,560 and 1,980 replayed runs and the earlier audits, we record every round at which a confidence interval fails to contain the true mean.
4. **AML application**. Certified search is applied to a BeatAML-calibrated AML network, which identifies a super-additive combination of ibrutinib, midostaurin and an ERK-inhibitor class in the model. We test its dependence on the calibration, the network structure, the drug-to-gene map and genotype.

### Algorithm 1

Certified search (as implemented).

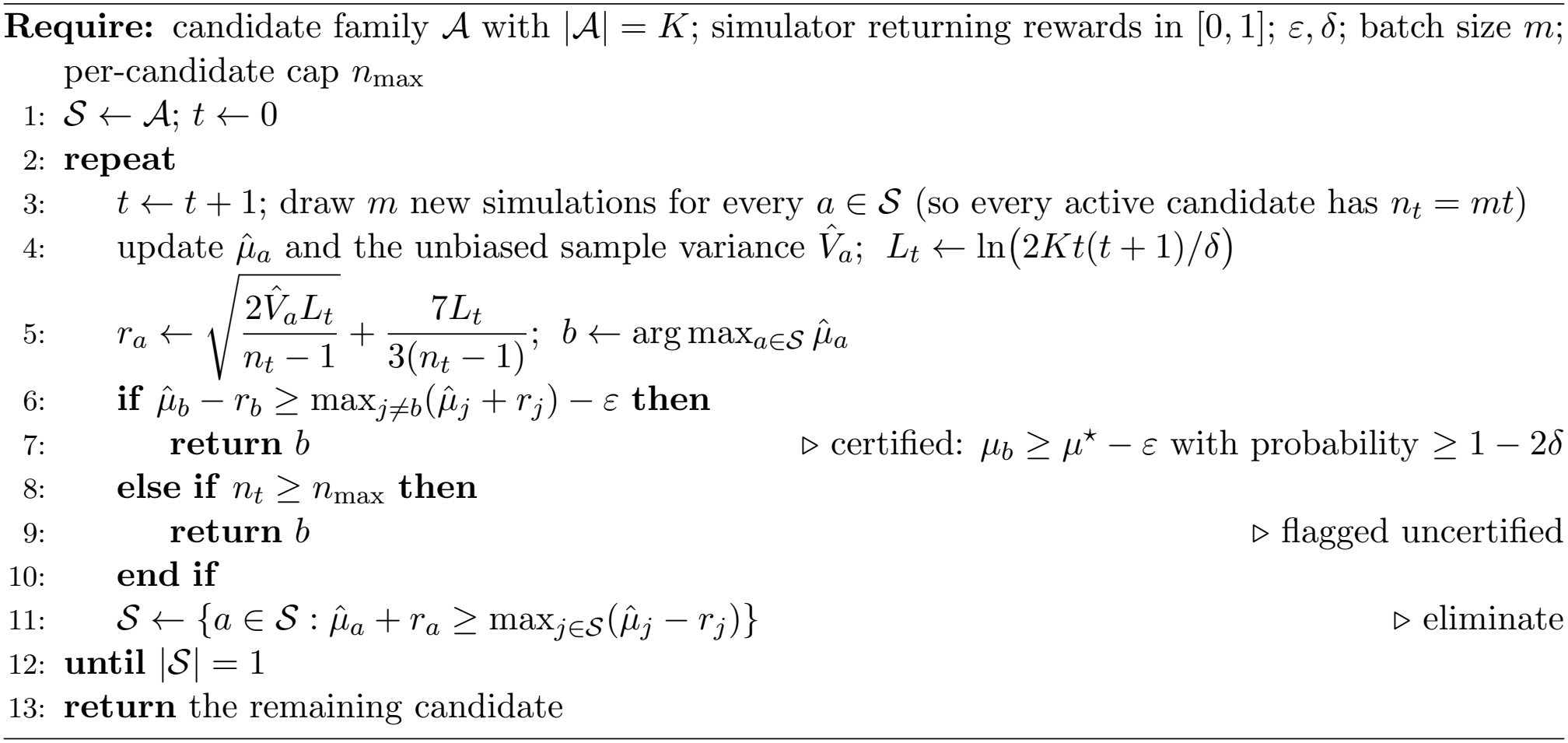

A single code file generates all numbers, tables and figures (Supplementary Note 7).

## 2 Results

### 2.1 Certified search

Let *A* be a finite family of interventions, for example all sets of at most three targetable genes clamped to zero. One stochastic simulation of intervention *a* returns a reward *X* ∈ [0, 1], here the fraction of output nodes that end in their desired state. We want *µ*_*a*_ = E[*X* | *a*].

Certified search works in rounds (Algorithm 1). Every active candidate gets the same number of new simulations, empirical-Bernstein confidence intervals [12, 13] are updated, and any candidate whose upper bound falls below the best lower bound is dropped. The search stops when the lower bound of the empirical best is within *ε* of every other candidate’s upper bound (Fig. 1a). If it stops this way, the returned set satisfies *µ*_*â*_ ≥ max_*a*_ *µ*_*a*_ *− ε* with probability at least 1 *−* 2*δ* (Theorem 1). A run that reaches its per-candidate sample cap first is reported as uncertified. The guarantee is about the model’s expected score over the candidate family. It says nothing about biology.

**Figure 1.**
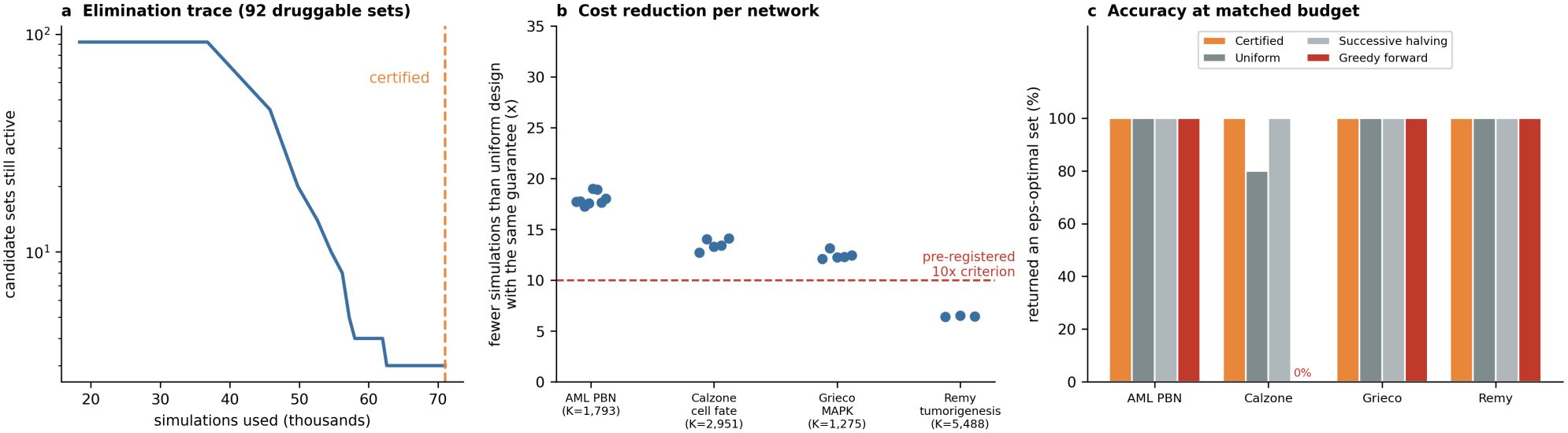
Certified search reaches a certificate at a fraction of the uniform cost. **a**, Elimination trace for the 92-set druggable AML search: candidate sets still active against simulations used. **b**, Reduction in simulations relative to the uniform design with the same guarantee, one point per run; the dashed line is the pre-specified 10*×* criterion. **c**, Percentage of runs that returned an *ε*-optimal set at matched budget.

Uniform sampling with the same guarantee needs ⌈ln(2*K/δ*)*/*(2*ε*^2^)⌉ simulations for each of *K* candidates [14]. Every reduction we quote is relative to this number, so it depends on *ε* and *δ*, as we show below.

#### Sampling rules

The certificate depends only on the confidence sequence and the stopping rule, not on which candidates are sampled. We therefore also consider LUCB [8] and UGapE [9], which sample the empirical best candidate and its strongest challenger one batch at a time, and a batch-parallel LUCB that samples the best candidate and the *B −* 1 strongest challengers in every round [11]. With *B* equal to the number of workers, a round of batch LUCB takes one batch time. The confidence sequence is indexed by the number of batches each candidate has received, so that Theorems 1 and 2 (Supplementary Note 1) hold for all of these rules.

### 2.2 Pre-specified benchmark

To see how the method behaves in practice, we ran a benchmark whose success criteria were fixed in advance. Before running it we wrote down the networks, objectives, dynamics, *ε* = 0.02, *δ* = 0.05, three baselines and three success criteria (Supplementary Table 7). The criteria were: (C1) an *ε*-optimal set in at least 95% of certified runs; (C2) at least ten times fewer simulations than the guaranteed-uniform design in every network; (C3) accuracy at matched budget at least as good as every baseline. “Pre-specified” here means time-stamped inside our project. We did not deposit the protocol in a public registry.

The benchmark used the BeatAML-calibrated 24-gene AML probabilistic Boolean network [15] and three published cancer Boolean models from the PyBoolNet repository [16]: cell-fate decision after death-receptor engagement [17], MAPK-driven cell-fate decision [18] and bladder tumorigenesis [19]. We had also planned to use a model of T-cell large granular lymphocyte leukemia [1], but its untreated score was already 0.925, so no intervention could be separated from any other and we dropped it (Methods). In all four benchmark networks the clampable set excludes the output nodes (Methods). For each network we screened every candidate and then re-estimated the top 30 with 60,000 simulations each to obtain ground truth.

Table 1 and Fig. 1b,c show the outcome. Every certified run returned an *ε*-optimal set (21/21), so C1 was met. That is a count of successes, though, and 21 of 21 does not establish a success probability of 95%. The exact one-sided 95% lower confidence bound is 86.7%. The saving was 17.3–19.0*×* on the AML network, 12.7–14.1*×* on Calzone and 12.1–13.1*×* on Grieco. On the largest family (Remy, 5,488 sets) it was only 6.4–6.5*×*, so **C2 was not met**. C3 held with ties: successive halving found an *ε*-optimal set in 21/21 runs, uniform allocation in 20/21 and greedy forward selection in 16/21. Greedy selection failed in every Calzone run, where the best set cannot be reached by adding one gene at a time.

**Table 1:** Benchmark summary. *K* is the number of candidate sets; “untreated” and “best” are the model scores of the empty set and of the best set. Success counts are runs that returned an *ε*-optimal set. Each baseline used the same number of simulations as the corresponding certified run. The AML network is scored with the strict five-marker reward (Methods).

| Network | *K* | Untreated | Best | Runs | Reduction | Uniform | Halving | Greedy |
|---|---|---|---|---|---|---|---|---|
| AML PBN (KGBN) | 833 | 0.445 | 0.684 | 8 | 17.3–19.0× | 8/8 | 8/8 | 8/8 |
| Calzone cell fate | 2,951 | 0.603 | 0.809 | 5 | 12.7–14.1× | 4/5 | 5/5 | 0/5 |
| Grieco MAPK | 1,275 | 0.717 | 0.877 | 5 | 12.1–13.1× | 5/5 | 5/5 | 5/5 |
| Remy tumorigenesis | 5,488 | 0.520 | 0.659 | 3 | 6.4–6.5× | 3/3 | 3/3 | 3/3 |

Three things limit how much these numbers show. At matched budget the simple baselines were usually right as well, so certified search adds a certificate and an adaptive stopping rule, not a higher hit rate on these networks. Repetitions per network (three for Remy, which we stopped for lack of time) are too few to estimate a miss probability. And ground truth comes from a screen followed by re-estimation of the top 30 sets, which could in principle miss a better set outside the top 30. The margin between the best refined estimate and the 30th screening score is large in every network (Supplementary Table 9); the union bound on missing a better set is at most 1.

### 2.3 Comparison with other methods and ablations

Matched-budget accuracy cannot separate procedures that usually find the right set, so we compared procedures by the number of simulations each needs to reach the same certificate. We replayed candidate families from simulated pools, so that every true mean is known exactly: the 92 druggable AML sets (92 sets; 100,000 simulations each), all 833 sets of up to three of the 17 clampable AML genes, the 15 highest-scoring of these, and all candidate sets of the three published networks (Grieco, 1,275 sets; Calzone, 2,951; Remy, 5,488), each with 40,000 simulations per set, the benchmark’s cap. Draws are independent samples from each candidate’s pooled reward distribution (Methods). Certified search was compared with LUCB [8] and UGapE [9], with three ablations: no elimination (the stopping rule alone), a Hoeffding radius, and checks at doubling sample sizes. All procedures use the same *δ* = 0.05, batch size, cap and, except for the Hoeffding ablation, the same empirical-Bernstein radius, with the union bound taken over (candidate, number of batches) so that the guarantee also covers adaptive sampling. They differ only in how simulations are allocated and when the search stops (Table 2, Fig. 2).

**Table 2:**
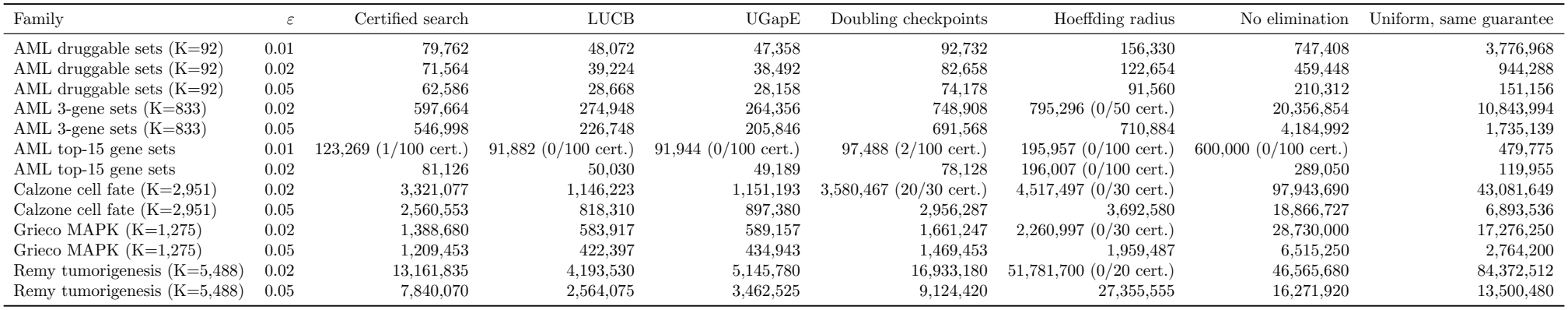
Mean simulations to certificate for certified search, two index methods and three ablations. All methods use the same *δ* = 0.05, batch size, per-candidate cap and (except the Hoeffding ablation) the same empirical-Bernstein radius; they differ in allocation or stopping only. Candidate rewards are replayed from simulated pools, so the true means are exact (Methods). Between 20 and 100 runs per cell (Supplementary Table 20). Where fewer than all runs certified before the cap, the count is given.

**Figure 2.**
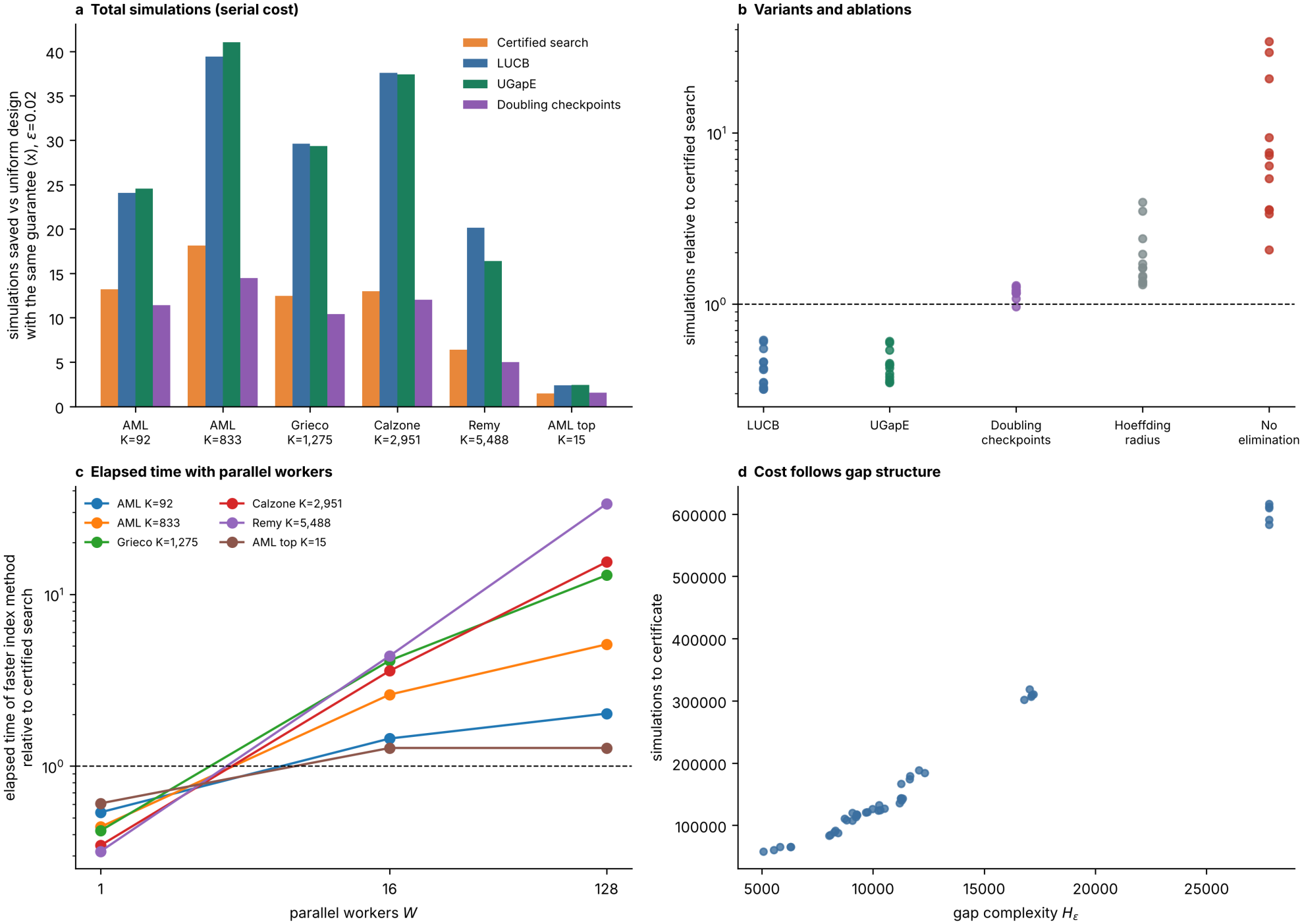
Allocation rules, ablations, elapsed time and gap structure. **a**, Saving relative to the uniform design with the same guarantee for certified search (successive elimination), LUCB, UGapE and the doubling-checkpoint variant, at *ε* = 0.02. **b**, Simulations of each variant relative to certified search, one point per setting in which certified search certified in at least 90% of runs. **c**, Elapsed time of the faster of LUCB and UGapE relative to certified search, against the number of parallel workers *W* (values above 1 favour certified search). **d**, Simulations to certificate against the gap complexity *H*_*ε*_ = ∑_*a*_ max(Δ_*a*_, *ε*)^*−*2^ in the controlled test of Supplementary Table 22.

Every one of the 4,560 runs returned an *ε*-optimal set (4,560/4,560) and in none did any confidence interval exclude the true mean (0 violations). The comparison supports four statements. (i) *Elimination is the main source of saving*. Without it, the stopping rule needs 2.1 to 34.1 times more simulations. (ii) *The variance-adaptive radius matters*. A Hoeffding radius needs 1.3 to 3.9 times more simulations and failed to certify before the cap in 330/760 runs across all settings (every family at *ε* = 0.02 except the 92 druggable sets, and the 15-set family at both *ε*), because the simulated scores have far less than the worst-case variance. (iii) *Index methods use fewer simulations in total but are sequential*. With the same radius, LUCB and UGapE needed 1.6 to 3.1 and 1.6 to 2.9 times fewer simulations than certified search, respectively. That gain comes with 50 to 45971 sequential decision rounds for UGapE, against 12 to 347 parallel rounds for certified search. Against the uniform design with the same guarantee, the saving at *K* = 92, *ε* = 0.01 was 47.4*×* for certified search, 78.6*×* for LUCB and 79.8*×* for UGapE; at *K* = 833, *ε* = 0.02 it was 18.1*×* for certified search and 41.0*×* for UGapE (means over replays; the single run quoted below for the druggable search saved 53.2*×*). *In elapsed time the order reverses once simulations can run in parallel*. With *W* workers a round of certified search takes ⌈*A/W* ⌉ batch times for *A* active candidates, whereas a LUCB or UGapE round is one batch time because the next choice depends on the previous result (Supplementary Table 23). Compared with the faster of the two index methods, certified search was 1.2 to 4.4 times faster with *W* = 16 and 1.3 to 34 times faster with *W* = 128, and 13 to 34 times faster with *W* = 128 on the three published networks (Fig. 2c). In serial (*W* = 1) the index methods remain 1.6 to 3.1 times cheaper. The model charges no dispatch latency, which favours the sequential methods. (iv) *Checkpoints trade a little cost for many fewer rounds*. Checking at doubling sample sizes cost 0.96 to 1.29 times as many simulations as certified search and reached the certificate in 5 to 9 rounds, which suits settings where each round is an expensive batch job. The batch size is not critical: the mean cost varied by at most 11% for *m* between 100 and 400 and rose by 1.31 to 1.52 times at *m* = 1,000 (Supplementary Table 21). The simple fixed-budget baselines also did well with the budget of each certified run: uniform allocation returned an *ε*-optimal set in 759/760 replays and successive halving in 760/760, so what certified search adds over them is the certificate and the stopping rule. The uniform design sized to the guarantee was correct in 760/760 replays at the stated cost. At the pre-registered *ε* = 0.02 all 330 replayed runs of certified search returned an *ε*-optimal set (exact one-sided 95% lower bound 99.1%), which complements the 21 runs of the benchmark itself.

(v) *Batch-parallel sampling removes the sequential bottleneck*. Batch LUCB with the same confidence sequence, stopping rule, batch size and cap, and *B* equal to the number of workers, was at most 1% slower than successive elimination in any of the 12 settings (1.01 to 3.2 times faster with *W* = 16, 0.99 to 3.1 times with *W* = 128; medians 2.4 and 1.3), and all 1,980/1,980 of its runs were *ε*-optimal with 0 interval violations (Table 3). With *B* = 16 it also used 2.0 to 3.1 times fewer simulations than successive elimination on the three published networks. On the two small families (92 and 15 sets) it used more simulations, because it samples *B* candidates per round whether or not they are still competitive, and its elapsed time was equal. The correctness of batch LUCB follows from the confidence sequence (Supplementary Theorem 2). We have no sample-complexity bound for it, and the batch-pull idea is due to Jun and colleagues [11]; what we add is its combination with the empirical-Bernstein certificate and its evaluation on simulation-based intervention search.

**Table 3:** Batch-parallel LUCB against successive elimination (SE) and sequential UGapE. Mean elapsed time with *W* = 16 workers (columns 3–5) and *W* = 128 workers (columns 6–8), in batch times, and mean total simulations (columns 9–11). Batch LUCB samples the empirical best candidate and the *B −* 1 strongest challengers in each round, with *B* = *W* for the elapsed-time columns and *B* = 16 for the total-simulations column; it uses the same confidence sequence, stopping rule, batch size and cap as the other columns. 20 to 100 runs per row.

| Family | ε | SE | Batch LUCB | UGapE | SE | Batch LUCB | UGapE | SE | Batch LUCB (*B* = 16) | UGapE |
|---|---|---|---|---|---|---|---|---|---|---|
| AML druggable sets (K=92) | 0.01 | 10,766 | 8,690 | 30,158 | 7,978 | 8,084 | 29,158 | 79,762 | 138,240 | 47,358 |
| AML druggable sets (K=92) | 0.02 | 8,032 | 6,188 | 21,292 | 5,256 | 5,166 | 20,292 | 71,564 | 98,208 | 38,492 |
| AML druggable sets (K=92) | 0.05 | 5,094 | 3,276 | 10,958 | 2,314 | 2,198 | 9,958 | 62,586 | 51,616 | 28,158 |
| AML 3-gene sets (K=833) | 0.02 | 55,770 | 32,732 | 186,356 | 27,538 | 25,584 | 181,756 | 597,664 | 522,212 | 264,356 |
| AML 3-gene sets (K=833) | 0.05 | 36,672 | 13,674 | 127,846 | 8,354 | 5,622 | 123,246 | 546,998 | 217,284 | 205,846 |
| AML top-15 gene sets | 0.02 | 19,131 | 18,919 | 47,789 | 19,131 | 19,359 | 47,789 | 81,126 | 283,785 | 49,189 |
| Calzone cell fate (K=2,951) | 0.02 | 225,133 | 82,197 | 874,593 | 51,330 | 37,340 | 858,493 | 3,321,077 | 1,300,983 | 1,151,193 |
| Calzone cell fate (K=2,951) | 0.05 | 163,097 | 51,243 | 620,780 | 23,060 | 10,387 | 604,680 | 2,560,553 | 818,993 | 897,380 |
| Grieco MAPK (K=1,275) | 0.02 | 95,630 | 43,833 | 469,657 | 29,950 | 24,167 | 462,657 | 1,388,680 | 700,833 | 589,157 |
| Grieco MAPK (K=1,275) | 0.05 | 78,017 | 25,673 | 315,443 | 12,310 | 6,417 | 308,443 | 1,209,453 | 410,273 | 434,943 |
| Remy tumorigenesis (K=5,488) | 0.02 | 826,450 | 272,510 | 4,631,280 | 106,800 | 34,405 | 4,601,280 | 13,161,835 | 4,184,090 | 5,145,780 |
| Remy tumorigenesis (K=5,488) | 0.05 | 491,330 | 163,675 | 2,948,025 | 62,625 | 21,405 | 2,918,025 | 7,840,070 | 2,545,920 | 3,462,525 |

The benchmark compares answers with ground truth, but it does not test the event that the proof relies on, namely that every confidence interval contains the true mean. We tested this directly on the 92-set druggable family. We estimated every set’s true score with 60,000 simulations (standard error about 0.001, ten times smaller than the radii in use), ran 60 independent certified searches (*ε* = 0.01, *δ* = 0.05), and checked at every round whether every active interval contained the true mean. All 60 runs certified and 60 returned an *ε*-optimal set. In 0 runs any interval was violated. The mean cost was 79,107 simulations (SD 5,357).

That family has one clear winner, so we repeated the audit on a harder one: the 15 AML gene sets with the highest screening scores, of which 2 lie within *ε* = 0.02 of the best. All 15 runs certified, 15 returned an *ε*-optimal set, and 0 violated an interval (Supplementary Table 19). These audits are still modest. Zero violations in 60 or 15 runs only puts the violation rate below 4.9% and 18.1% respectively (one-sided 95% bounds). They are consistent with the nominal level of 2*δ* but do not demonstrate it, and the bound looks conservative, as a union bound over all candidates and rounds should.

The size of the saving depends strongly on the accuracy target. On the druggable family the mean reduction was 180.8*×* at *ε* = 0.005, 48.8*×* at 0.01, 13.5*×* at 0.02 and only 2.5*×* at 0.05 (Fig. 3b;

**Figure 3.**
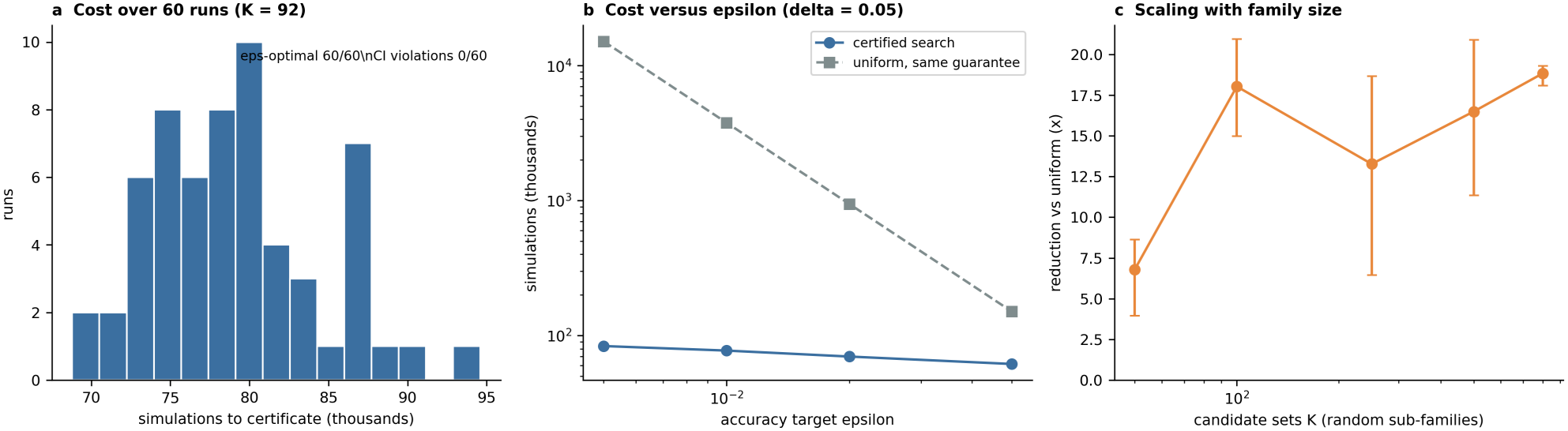
Cost of certification. **a**, Simulations to certificate over 60 independent runs on the 92-set druggable family; annotations give the audit counts. **b**, Mean cost against the accuracy target *ε* for certified search and for the uniform design with the same guarantee (*δ* = 0.05). **c**, Reduction against candidate-family size for random sub-families of the AML gene sets (*ε* = 0.02; bars show the range over three sub-families).

Supplementary Table 13). The reason is that eliminating the clearly poor candidates costs the same whatever *ε* is (about 61,700 simulations here), while the uniform design grows as 1*/ε*^2^. Certified search therefore pays off when *ε* is small compared with the score differences that matter, and it gives almost nothing at coarse tolerances. We saw a smooth decline and no sharp transition. The dependence on *δ* is weak: 52.0*×* at *δ* = 0.01 and 45.6*×* at *δ* = 0.2 for *ε* = 0.01.

The variance-adaptive radius made certification much cheaper in our tests. When we replaced it with a Hoeffding radius (range 1), the same search at *ε* = 0.02 gave a mean reduction of 7.6*×* instead of 13.5*×* and needed 1.8 times more simulations; 0 of three Hoeffding runs failed to certify within the sample cap. Simulated scores in these networks have small variance compared with the worst case of 1*/*4, and the empirical-Bernstein bound takes advantage of that. The cost of the stricter constant that would give a proven confidence of 1 *− δ* instead of 1 *−* 2*δ* (replacing 2 by 4 inside the logarithm) was 5.0% more simulations in 20 paired runs.

Next we asked how the cost changes with the size of the candidate family and with simulation noise. We drew random sub-families of the AML gene sets with *K* from 50 to 800 (three per size, *ε* = 0.02). Every run certified (15/15), with reductions of 4.0–20.9*×*. The reduction did not rise or fall steadily with *K*: it averaged 6.8*×* at *K* = 50, 18.0*×* at *K* = 100 and 18.8*×* at *K* = 800 (Fig. 3c; Supplementary Table 14). Over the same range the mean number of near-ties (sets within 0.04 of the best in a screen) went from 3.7 to 2.7. Random sub-families confound size with gap structure, so we ran a controlled test (Supplementary Table 22; Fig. 2d). At *K* = 100, including the *j* best sets with *j* = 1, 3, 5, 10, 20 raised the cost from 63,010 to 141,153 simulations and lowered the reduction relative to uniform sampling from 16.5*×* to 7.3*×*. At *j* = 5 near-ties, increasing *K* from 50 to 833 raised the cost from 87,517 to 602,530 simulations but raised the reduction from 5.4*×* to 18.0*×*, since the uniform design grows faster with *K*. Across all runs the cost correlated with the gap complexity *H*_*ε*_ = ∑_*a*_ max(Δ_*a*_, *ε*)^*−*2^ (Spearman *ρ* = 0.98). Cost therefore follows the gap structure, with a further logarithmic dependence on *K*, and the reduction is largest for large families with few near-ties, which is consistent with the smaller saving on the Remy network (many near-ties, not tested directly).

### 2.4 Application to AML

We then applied the method to AML, using a 24-gene probabilistic Boolean network whose rule probabilities were fitted to BeatAML drug-response data for five drugs [15, 20]. Our first analysis scored interventions with a seven-marker reward (TP53 and CDKN2A on; BCL2, MYC, CCND1, AKT1 and STAT5A off) and allowed every gene to be clamped, including the reward genes. Two of the drug targets, BCL2 (venetoclax) and AKT1 (AKT inhibitors), were therefore reward nodes, and clamping them satisfied part of the reward by construction. We found the problem while reviewing an earlier version of the analysis. The consequence is large here: no gene’s rule uses BCL2 as a regulator (0 rules), so venetoclax cannot change any other node in this network. In the first analysis venetoclax gained 10.9 points over untreated. Under the corrected policy it gains 0.0 (Fig. 4; Supplementary Table 5). The first analysis’ headline pair (midostaurin + venetoclax, +19.1 points) falls to +1.5 points, which is the gain from midostaurin alone (+1.5), and its best three-gene set (BCL2 + MYC + SYK) was mostly the direct clamping of two output nodes.

**Figure 4.**
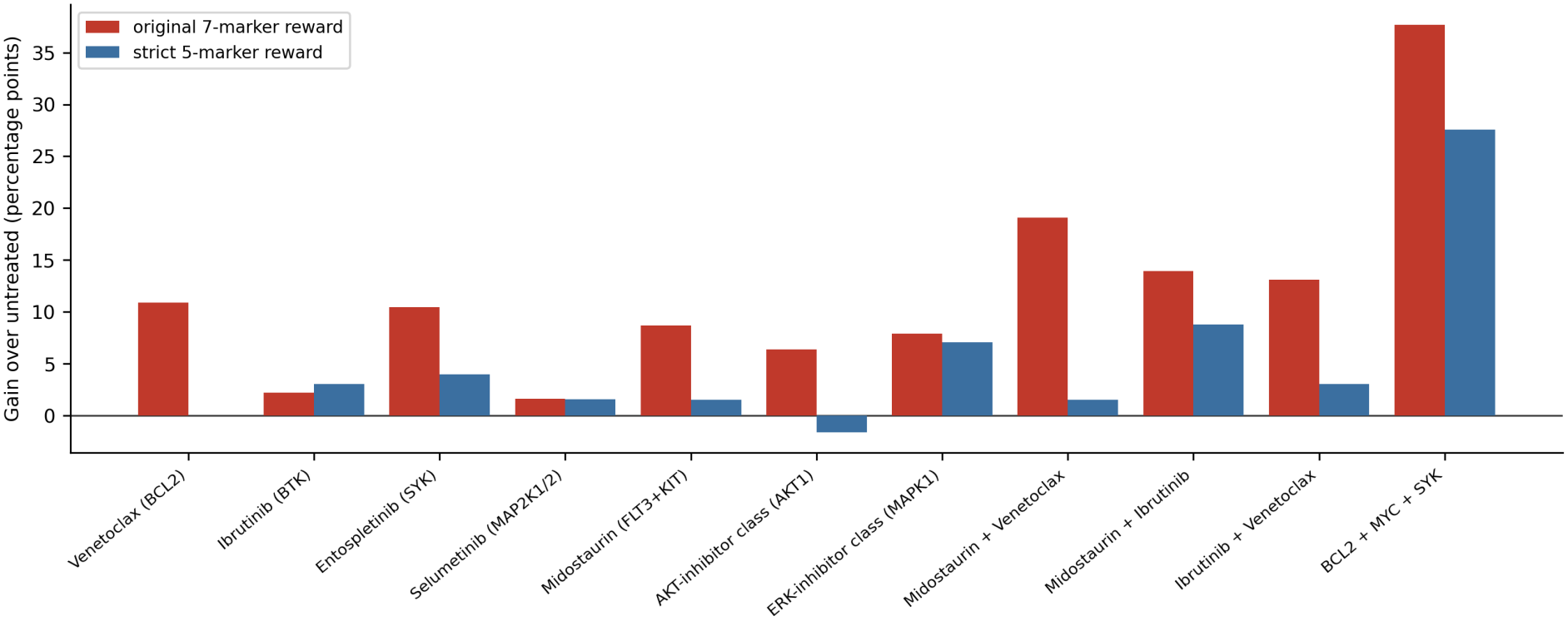
Effect of the clamping policy. Gain over the untreated score for the same simulations scored with the original seven-marker reward (red; includes the drug targets BCL2 and AKT1) and the strict five-marker reward used in the main analysis (blue). Venetoclax has no effect under the strict reward.

All AML results in this paper use the strict policy: TP53 and CDKN2A (desired on) and MYC, CCND1 and STAT5A (desired off) form the reward; none of them can be clamped; BCL2 and AKT1 remain clampable drug targets but are not part of the reward, so a drug acts only through network dynamics. We regard this as the definitive policy because a reward that an intervention can set directly measures access to the reward nodes and not the effect of the intervention on the network; under the strict policy every gain has to propagate through the network’s rules, which is what a drug target does. The first analysis is superseded and is kept only in Supplementary Table 5. The reward is the mean of five marker indicators.

Under this policy, among the five drugs and five selected pairs, midostaurin + ibrutinib scored highest (53.4% against 44.7% untreated; Fig. 5a). It is super-additive under both a linear-additive null and a marker-level Bliss-independence null (Methods): the observed score exceeds the additive expectation by 4.21 percentage points (SD 0.19 over 30 replicate batches) and the Bliss expectation by 4.54 points, and it beats the best single agent by 5.7 points (Supplementary Table 6). The effect is modest. Cohen’s *d* against the nearest competitors is at least 0.21 (probability of superiority at least 0.56), and midostaurin + entospletinib is sub-additive (-1.3 points), which is consistent with the two clamps acting on a shared route in the network. Among all 680 three-gene sets of the 17 clampable genes the pair sits at percentile 92.8 (rank 50), and the best three-gene set, BTK+FLT3+MAPK1, reaches 68.5%, so, within the model, the pair is the highest-scoring of the drug pairs we considered, and it is not the highest-scoring three-gene target set.

**Figure 5.**
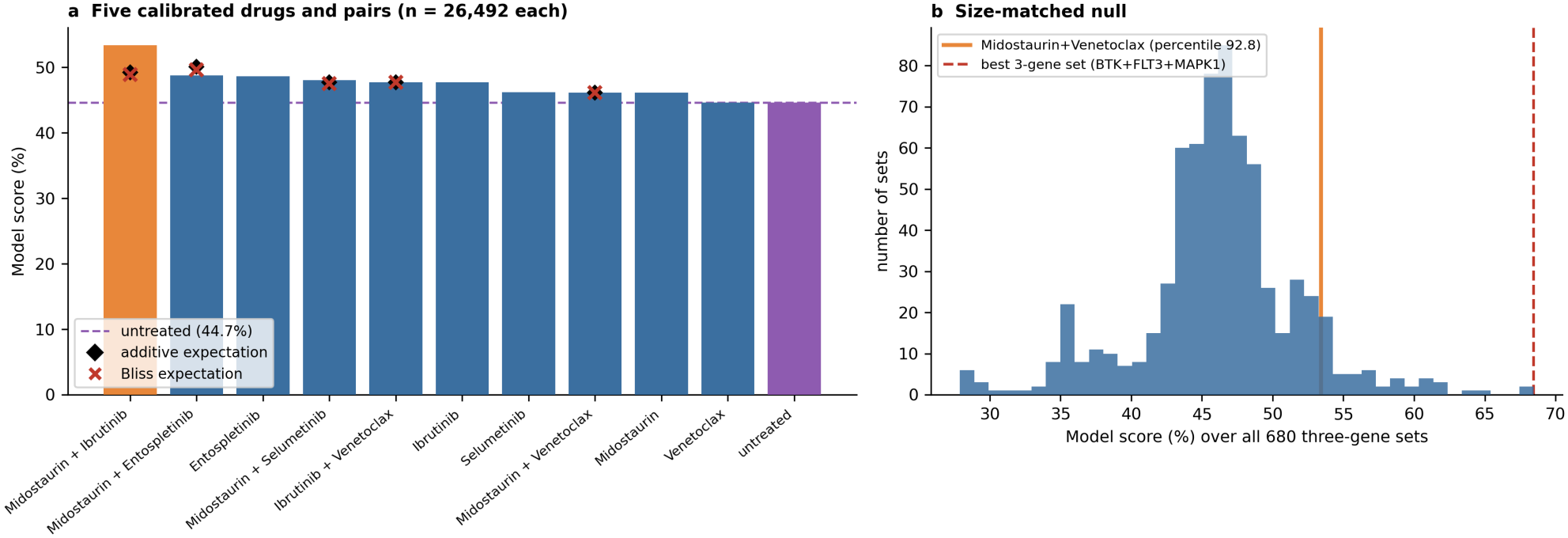
Five calibrated drugs and pairs under the strict reward. **a**, Model score of untreated, single drugs and pairs (*n* = 26,492 simulations each; diamonds mark the additive expectation and crosses the Bliss expectation). **b**, Scores of all 680 three-gene clamp sets; the pair clamps three genes.

The ranking was stable in the tests we ran. In 200 independent simulation batches the pair ranked first in 100% at the original sample size (*n* = 300) and in 100% at *n* = 1,000. With mutation status held fixed it ranks first in all eight FLT3/NPM1/DNMT3A genotypes (Fig. 6a), with a gain over untreated of 9.5 points in the FLT3-, NPM1- and DNMT3A-mutant stratum and 6.5 points in the corresponding FLT3-wild-type stratum. Across 15 distinct calibrations shipped with the KGBN toolkit it ranks first in 14 and second in 1 (mean Kendall *τ* = 0.68), and with rule-probability noise at *κ* = 10 it stays first in 96% of 100 perturbations (Fig. 6b,c). A FLT3-only inhibitor (gilteritinib-like) combined with ibrutinib scores 53.32%, against 53.47% for midostaurin (FLT3 and KIT), so within the model the KIT arm adds almost nothing. The calibration is weaker for midostaurin than for venetoclax: the pooled correlation between predicted and observed BeatAML response is 0.879, but the correlation within a single drug across the eight genotypes is 0.94 for venetoclax and only 0.29 for midostaurin.

**Figure 6.**
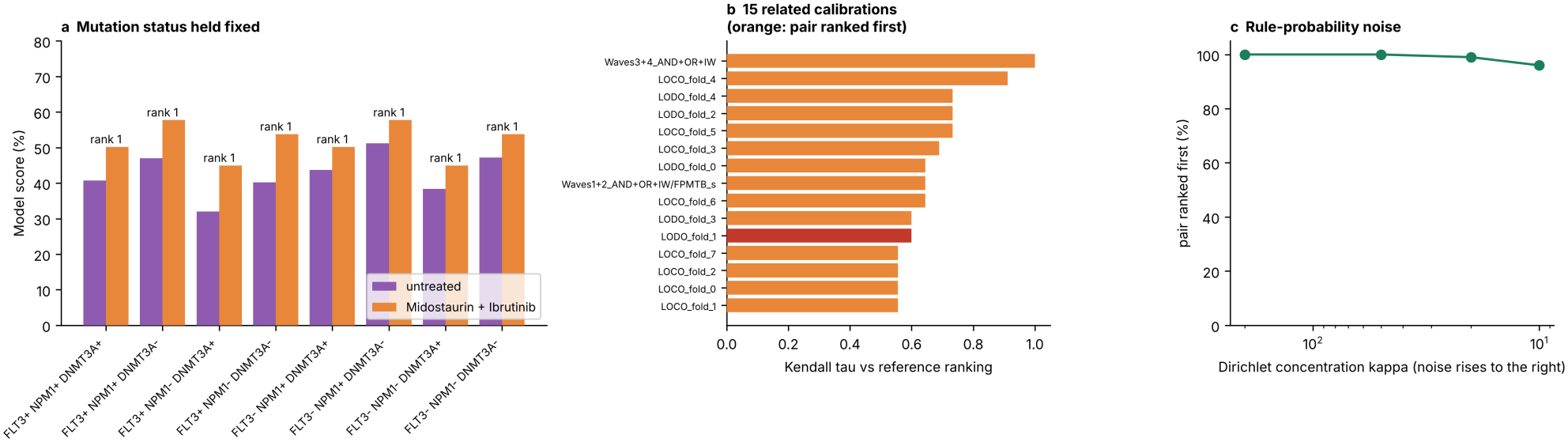
Genotype dependence and robustness of the pair ranking. **a**, Mutation status held fixed at every step for all eight FLT3/NPM1/DNMT3A genotypes; labels give the pair’s rank. **b**, Kendall rank agreement with the reference ranking across 15 distinct KGBN calibrations (orange: pair ranked first). **c**, How often the pair is ranked first under Dirichlet noise on every rule-probability vector (small *κ* means more noise).

The pairs above came from a short list we selected ourselves, so we also asked which drug combination the model ranks highest. For this we restricted the family to sets of up to three drugs drawn from eight units with clear gene targets: venetoclax (BCL2), ibrutinib (BTK), entospletinib (SYK), selumetinib (MAP2K1 and MAP2K2), midostaurin (FLT3 and KIT), a gilteritinib-class FLT3 inhibitor (FLT3), an AKT-inhibitor class (AKT1) and an ERK-inhibitor class (MAPK1). This gives *K* = 92 candidate sets. We set *ε* = 0.01 and *δ* = 0.05. The search certified its answer with 71,000 simulations against 3,776,968 for the uniform design (53.2*×* fewer; Fig. 1a) and returned **ibrutinib + midostaurin + an ERK-inhibitor class**. Its score was 0.710, against 0.448 untreated. The additive expectation was 0.560 (excess 0.149) and the Bliss expectation 0.551 (excess 0.158; Fig. 7a; Supplementary Table 10).

**Figure 7.**
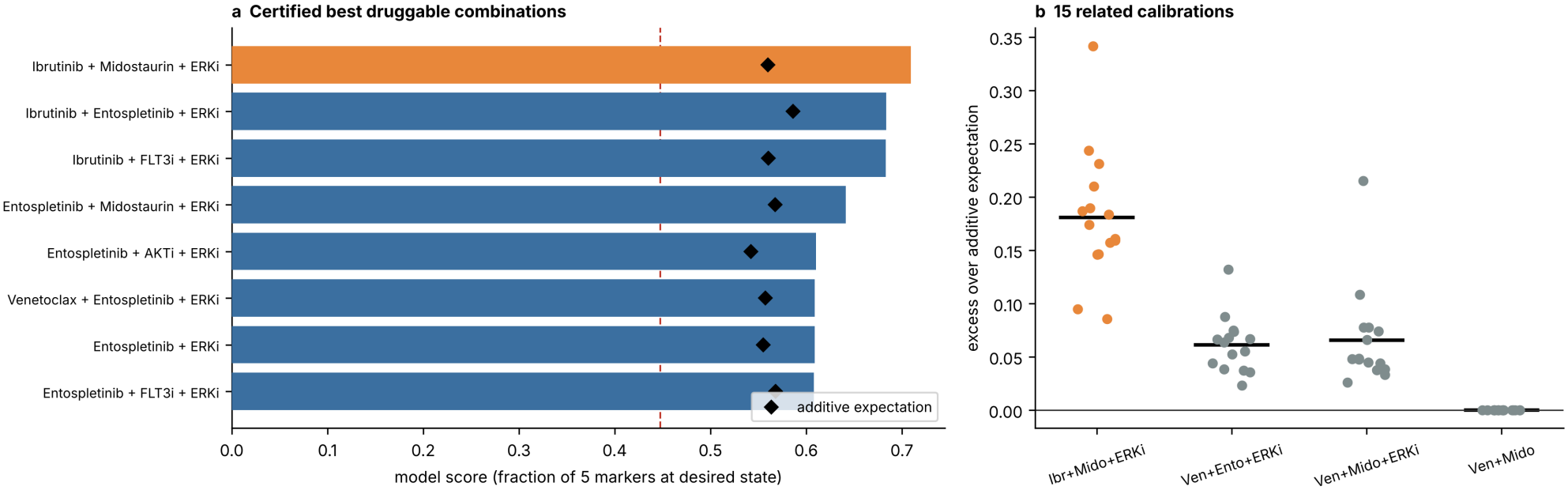
In the model, a super-additive triplet is the certified best druggable combination. **a**, The eight best druggable combinations by model score; diamonds mark the additive expectation and the dashed line the untreated score. **b**, Excess over the additive expectation for four sets in each of 15 distinct calibrations; bars are means.

The 15 calibrations shipped with the KGBN toolkit are distinct but not independent. They have identical rule sets and differ only in rule probabilities, which are fitted on overlapping data (leave-one-cohort-out and leave-one-drug-out folds, and different BeatAML waves). The pairwise Spearman correlation of their rule-probability vectors is 0.67 on average (minimum 0.36), and 0.81 (minimum 0.21) for the scores of the ten five-drug candidates. We therefore treat them as an ensemble of related fits and not as replicates. In our first analysis the triplet was compared with only three other sets in each calibration. We now ran the full 92-set certified search inside every calibration. The triplet was the certified best set in 15/15 of them (rank at most 3 in 15/15). Its excess over the additive expectation was positive in 15/15 (mean 0.182, range 0.083 to 0.342) and its excess over the Bliss expectation in 15/15 (mean 0.201, range 0.107 to 0.356; Supplementary Table 11).

Because the calibrations share their rule structure, they could all be wrong in the same way, so we also perturbed the structure directly (Fig. 8b; Supplementary Table 12). Removing the MAPK1 input to STAT5A left the excess at 0.143 (reference network: 0.152), and the triplet remained the best of the four triplets that can be formed from ibrutinib, midostaurin, entospletinib and the ERK-inhibitor class. Swapping the probabilities of the two main STAT5A rules gave an excess of 0.089. Across 60 random rewirings of one regulator the excess was positive in 100% of networks (mean 0.141) and the triplet was best in 100%; with three simultaneous rewirings the figures were 100% positive (mean 0.132) and 98% best. This is evidence of robustness to small structural changes in this network, not to uncertainty about the biology.

**Figure 8.**
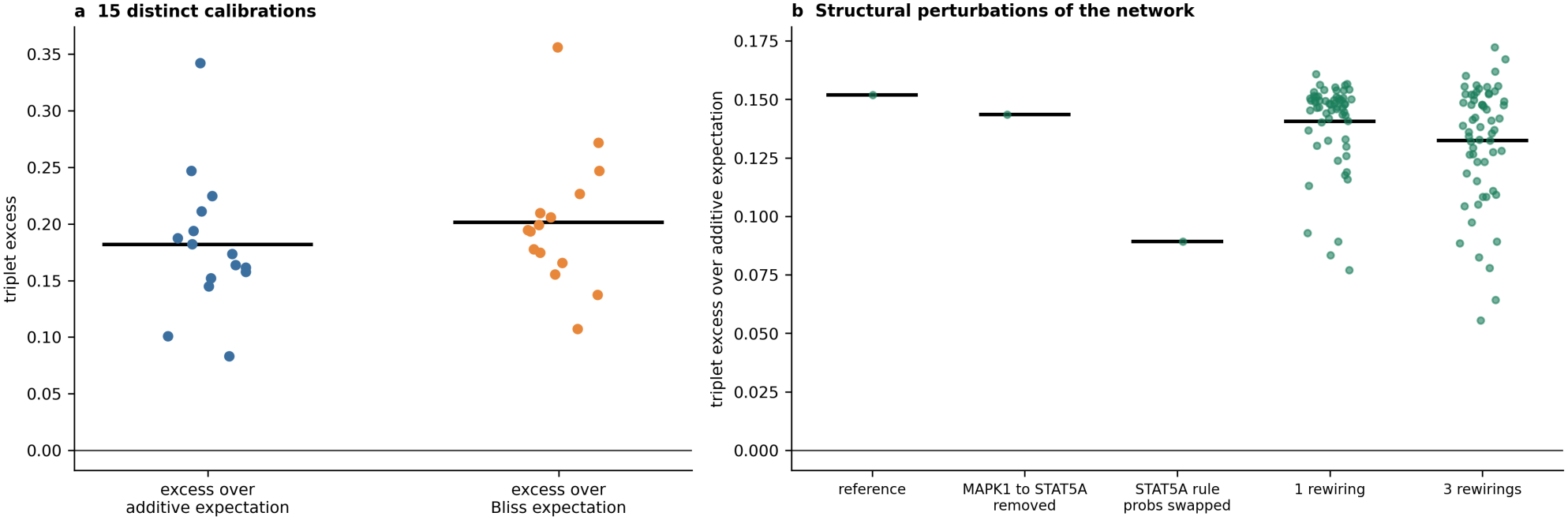
Robustness of the triplet’s super-additivity. **a**, Excess over the additive and over the Bliss expectation in each of 15 distinct calibrations (bars are means). **b**, Excess over the additive expectation in the reference network, after removing the MAPK1 input to STAT5A, after swapping the two main STAT5A rule probabilities, and under random rewiring of one or three regulators.

**Figure 9.**
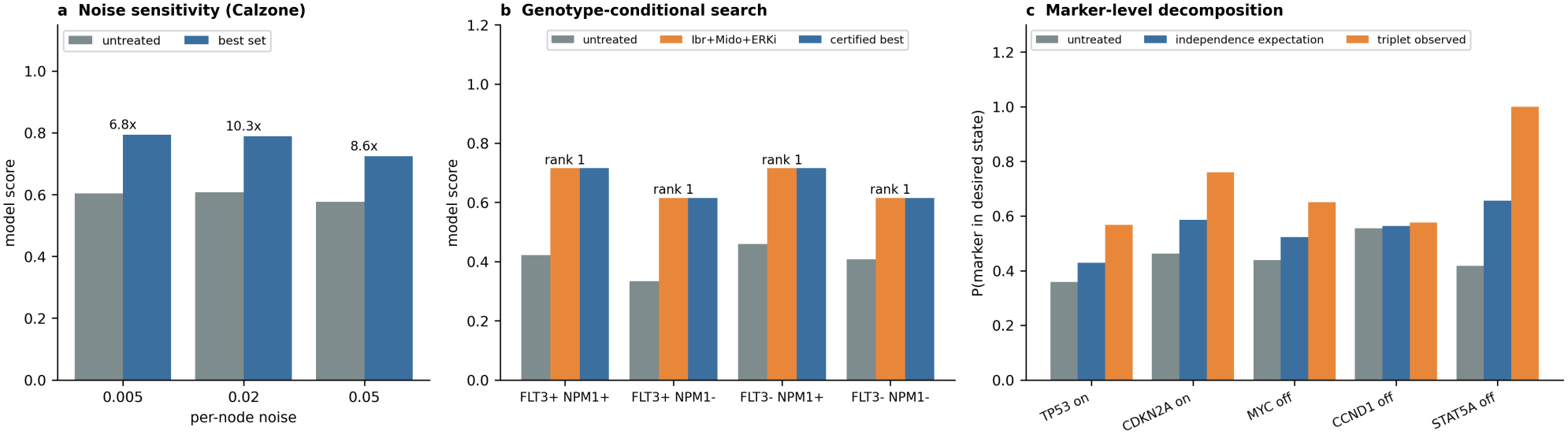
Noise, genotype and mechanism. **a**, Calzone network at three noise levels: untreated and best-set scores, with the reduction achieved by certified search written above each pair of bars. **b**, Genotype-conditional certified search: untreated score, the ibrutinib + midostaurin + ERK-inhibitor triplet and the certified best set in four FLT3*×*NPM1 strata (label: rank of the triplet). **c**, Probability that each output marker is in its desired state: untreated, independence expectation, and observed value for the triplet.

Several limits apply to this result. The ERK-inhibitor unit clamps MAPK1, which none of the five calibrated drugs target, so that part of the prediction lies outside the data the model was fitted to. The mapping from drug classes to genes is our assumption and needs review by a pharmacologist. Midostaurin is the drug for which the calibration is weakest. Super-additivity inside a Boolean model is a statement about parallel pathway redundancy and does not by itself show drug synergy, and the two nulls we use (linear additivity and Bliss independence of marker-level effects) are model-based, so the excess should not be read as a measure of drug synergy. The triplet is a model-derived result only; we have not compared it with any external data and draw no biological conclusion from it.

We also asked whether the result needs the real drug-to-gene map or arises for arbitrary maps (Fig. S4a,b; Table 25; Supplementary Note 9). This analysis was added after the main analyses and was not pre-specified. We kept the eight drug units and their sizes, gave each unit random targets among the 17 clampable genes, and ran the same 92-set search on 1000 decoy maps with 1,000 simulations per set; the real map was evaluated 30 times to show the noise of the procedure. We computed four statistics on the same decoy maps and report all of them, with a Holm adjustment over the four. The most conservative is the largest excess over additivity among all 92 sets of a map, because every decoy map also benefits from selecting its own best set: the real map gave 0.152 against a decoy median of 0.032 for the best-scoring set, and the raw *p* for the selection-aware statistic was 0.043 (Holm 0.070). For the other three statistics the raw *p* was 0.006 for the excess over additivity of the best-scoring set (Holm 0.020), 0.005 for the same excess over Bliss independence (Holm 0.020), and 0.035 for the score of the best-scoring set (Holm 0.070). These statistics are not independent: in the real map the best-scoring set is also the set with the largest excess, so they differ in the strictness of the comparison and not in what was found. The raw *p* also moves with the noise in the real map’s own estimate (from 0.014 to 0.158 for the selection-aware statistic across single replicates). We treat the evidence as exploratory and modest, and we draw no conclusion from whether any single *p*-value falls below 0.05. Holding each of 17 genes off in turn (Fig. S4c; Supplementary Table 26), the triplet stayed the best set in 9 cases and among the top three in 15, and its excess stayed positive in all 17 (smallest 0.036). The two largest drops, when BTK or MAPK1 was held off, remove targets of the triplet itself, so a lower rank is expected there.

Because the case study rests on a fitted network, we checked how well that network predicts data it was not fitted to, using only held-out evaluations that ship with the KGBN toolkit (Methods; Supplementary Note 8, Fig. S3 and Table 24). The check concerns single-drug response and uses no combination data. Fitted on one BeatAML cohort and evaluated on the other, the network tracked the measured values with Spearman *ρ* = 0.65 and 0.69, against 0.77 for the measured training-cohort values themselves; against the independent FPMTB cohort (a different assay, *n* = 22, sign reversed because it scores sensitivity) it gave 0.60 and 0.56 against 0.61. This agreement is probably carried largely by differences between drugs, which the training data already contain; the within-drug results below are less consistent. For finer questions the evidence is weaker. With one drug left out of the fit, the pooled correlation was 0.28 (bootstrap 95% interval *−*0.05 to 0.59) against 0.11 for a trivial baseline, and with one mutation profile left out it was 0.28 against 0.37 for a drug-mean baseline. For ibrutinib and midostaurin, the two drugs of the pair, the held-out-drug correlations were 0.00 and 0.37 (permutation *p* = 1.00 and 0.36, *n* = 8 each), and the ordering of genotypes within a drug was not consistent between cohorts (ibrutinib *−*0.02 and 0.88; midostaurin 0.66 and 0.23). The triplet was the certified best set in all 13 calibrations that were fitted with a drug or a genotype held out, which fits the ranking being set mainly by the rule structure the calibrations share, so agreement among calibrations should not be read as evidence of predictive accuracy. These samples are small (*n* = 5 or 8 per fold), and the check says nothing about combinations. We therefore claim only that the network reproduces between-drug differences across cohorts, and we make no claim that it predicts the effect of the triplet.

We also asked whether the answer depends on genotype. We repeated the certified druggable search with mutation status held fixed in four FLT3*×*NPM1 strata (drug clamps take precedence; DNMT3A left unselected). Every stratum certified. In the FLT3-mutant, NPM1-mutant stratum the triplet scored 0.715 against 0.422 untreated (excess over additivity 0.235, over Bliss 0.240); in the FLT3-mutant, NPM1-wild-type stratum, 0.615 against 0.333 (0.169 and 0.177). In the two FLT3-wild-type strata it scored 0.715 against 0.460 and 0.615 against 0.408 untreated (excess over additivity 0.063 and 0.041; over Bliss 0.075 and 0.051). The triplet was the certified best set in all four strata (ranks 1, 1, 1 and 1), so the search did not return a genotype-specific winner. The treated scores are the same for the same NPM1 status because midostaurin clamps FLT3 whatever its mutation status; the strata differ in their untreated scores. What differs is the size of the super-additive excess, which is larger in the FLT3-mutant strata than in the wild-type strata (Supplementary Table 16).

To look at mechanism, we decomposed the excess by output marker, comparing the triplet with the marker-level independence expectation (Methods). The excess is spread over several markers and is not confined to one (Fig. 9c; Supplementary Table 17). In the reference network STAT5A has the largest share (it is off with probability 1.000 under the triplet against 0.656 expected; excess 0.344), but TP53, CDKN2A and MYC contribute as well (0.140, 0.174 and 0.128), and CCND1 almost nothing (0.013). The fitted STAT5A rules suggest one contributor: the dominant rule (probability 0.63) is BTK | FLT3 | MAPK1, so STAT5A is guaranteed off only when all three inputs are blocked. That is not the whole explanation. When the MAPK1 input to STAT5A is removed, the overall excess barely changes (see above), and when both the BTK and MAPK1 inputs are removed the STAT5A excess vanishes while the other markers keep an excess (Supplementary Table 18). We therefore do not claim that the STAT5A rule explains the triplet. What we can say is that, in this fitted model, the three components act on partly overlapping routes to several outputs, and that the rule structure around STAT5A contributes a part of it.

Two further analyses are reported in the Supplement, because they are only loosely tied to certified search. The first concerns uncertainty for individual patients in real per-patient BeatAML data. A bootstrap interval for a mean covers only about 9.2% of held-out patients, as expected, but a plain Gaussian prediction interval does as well as conformal prediction (92.5% and 90.2% in a leave-one-profile-out design), and all valid intervals are very wide (Supplementary Note 3). The second is a topological ranking score that we explored earlier in this project. It did not hold up: it correlated positively with outcome, opposite to our stated interpretation, and added little beyond trivial trajectory statistics (Supplementary Note 4).

## 3 Discussion

Certified search addresses one gap: model-based target rankings are usually reported without a statement of how confident one can be that the top set is the best in the model. On four networks it returned the correct set in every run, with a certificate, using between 6.4 and 19.0*×* fewer simulations than uniform sampling with the same guarantee. The ingredients are classical. What the work adds is their assembly for bounded simulation scores, an evaluation that attributes the saving to specific design choices, and audits of the guarantee.

### What the comparison shows

The saving over uniform sampling comes from two choices: eliminating candidates that are clearly worse (2.1 to 34.1 times fewer simulations than the stopping rule alone) and using a variance-adaptive radius (1.3 to 3.9 times fewer than Hoeffding). The allocation rule involves a trade-off, and it is the same on all four networks. With the same radius, UGapE and LUCB need 1.6 to 3.1 times fewer simulations in total than successive elimination, but each of their decisions depends on the previous one. Successive elimination evaluates all active candidates in parallel batches, so on 16 to 128 workers it reaches the certificate 1.2 to 34 times sooner, and by 13 to 34 times on the three published networks at 128 workers. Its doubling-checkpoint variant needs only 5 to 9 rounds. Sampling the index methods in parallel batches removes most of that disadvantage: batch LUCB was at most 1% slower than successive elimination and was up to 3.2 times faster. Which procedure to use depends on whether total compute or elapsed time is the scarcer resource. The saving is larger when *ε* is small relative to the gaps that matter, and it grows with family size at fixed near-ties, while many near-ties raise the cost. At matched budget, uniform allocation and successive halving were nearly always correct as well, so the practical addition of certified search is the stated confidence together with a stopping rule that spends simulations where the data require them.

### Practical guidance

Certified search suits cases where the objective is a bounded expectation of a stochastic simulation, the candidate family can be listed, and a statement of confidence in the winner is needed. The accuracy target *ε* should be chosen relative to the score differences that would change a decision. A variance-adaptive radius is worth using. It also pays to look at headroom first, since a network whose untreated score is already close to its best (T-LGL, 0.925) cannot be certified, and to check that no intervention target is also part of the reward. The certificate should be reported together with *ε, δ*, the number of simulations and whether the run hit its cap. The sampling rule should follow the hardware, and the certificate is the same for all of them. On a single processor, UGapE or LUCB need the fewest simulations. On a cluster, batch LUCB with the batch size set to the number of workers was the fastest or tied in every setting we tested. Successive elimination is a simpler alternative with a fixed round structure and a proven cost bound, and for families of about a hundred candidates or fewer it needs fewer simulations than batch LUCB at the same elapsed time.

### The AML case study

Under the strict reward, in which the intervention does not set any reward node directly, a pair (midostaurin + ibrutinib) comes out best among the five calibrated drugs, vene-toclax has no effect because no rule in the network uses BCL2 as a regulator, and a search over eight drug classes returns a triplet that includes an ERK-inhibitor class. The triplet is the best set in every related calibration, exceeds two model-based nulls, and keeps a positive excess under all structural perturbations and single-gene loss-of-function tests we ran. The decoy-map test gave modest evidence that a set this super-additive is unusual under random drug-to-gene maps. The result is a hypothesis generated by the model: it concerns the model’s expected score, and it has not been compared with combination data. The earlier version of this analysis allowed reward genes to be clamped; the corrected policy changes the venetoclax findings entirely, which illustrates that a certificate holds for the stated objective and is only as informative as that objective.

### Outlook

As stochastic models become larger and are used for more decisions, the number of simulations needed to separate candidates will matter more, and a stated confidence level makes a ranking easier to check and to reproduce. Natural next steps are certified search on other stochastic models and objectives, combination data to test the AML prediction, variation of the reward definition, horizon and drug-to-gene map, and comparison with further best-arm algorithms.

### Limitations

#### Scope of the guarantee

Everything concerns the *model’s* expected score; the AML predictions have not been compared with external combination data. The proven confidence level of the implemented procedure is 1 *−* 2*δ* (90% for *δ* = 0.05); the stricter constant costs 5.0% more. The sample-complexity bound of Theorem 3 holds with probability 1*−*3*δ* and is 7 to 31 times larger than the simulations used, so it supports the form of the cost but not its size. For LUCB and UGapE the comparison uses the same radius and a union bound over (candidate, number of batches); we did not tune either algorithm. The reduction factors are measured against a conservative guaranteed-uniform design and depend on *ε*. The benchmark itself, with 3 to 8 runs per network, used successive elimination only; the batch variant was evaluated on replayed pools, and we have a correctness proof but no sample-complexity bound for it. The elapsed-time model assumes that every batch takes the same time, that up to *W* batches run at once and that dispatch is free; real clusters add queueing and latency, which would favour the batch-parallel method further, and heterogeneous run times would weaken it.

#### Benchmark

The benchmark has three to eight repetitions per network, four networks, one noise level and one objective per network. One pre-specified criterion (ten-fold saving in every network) failed and one candidate network (T-LGL) was dropped for lack of headroom. Ground truth for the benchmark networks could in principle miss a better set outside the re-estimated top 30. We considered only inhibitory interventions. The method comparison replays pools drawn from each network, so the true means are exact; it covers the benchmark’s networks, one objective each, and the replayed runs of a family share its pool and are not independent of one another beyond it. The controlled test varies near-ties through the best-ranked sets in one family. The method comparison includes a family with many near-ties (Remy), but near-ties are varied systematically only in the controlled test. Our novelty check was a limited literature search.

#### AML model

The network is a 24-gene abstraction whose calibrations are related, share rule structure and are weak for midostaurin. On held-out public data it reproduced between-drug differences but was not shown to predict held-out drugs or genotypes better than simple baselines (Supplementary Note 8). The ERK-inhibitor unit lies outside the calibration, the drug-to-gene map is our assumption, super-additivity is judged against model-based nulls only, the mechanistic analysis is partial, and the reward definition, horizon and drug-to-gene map were not varied widely.

## 4 Methods

### Models and dynamics

The AML network is the 24-gene probabilistic Boolean network of the KGBN toolkit calibrated on BeatAML Waves 3+4 data [15]. At every synchronous step each gene independently samples one of its candidate rules. The published Boolean models come from the PyBoolNet repository [16]. We simulated them with synchronous updating in which each node’s update is flipped independently with probability 0.02 per step (a Boolean network with perturbations), for 50 steps from uniformly random initial states. For the AML network the horizon was 40 steps, with FLT3 and NPM1 set on at *t* = 0.

### Interventions, clamping policy and reward

An intervention clamps a set of nodes to zero after every update. For the three published Boolean models the clampable set excludes the output nodes, and the reward is the fraction of output indicators in the desired state at the final step (Supplementary Table 7). For the AML network we use a strict policy: the reward consists of TP53 and CDKN2A (desired on) and MYC, CCND1 and STAT5A (desired off); none of these five genes can be clamped, nor can the mutation nodes NPM1 and DNMT3A, so 17 genes are clampable; BCL2 and AKT1 are clampable and are not part of the reward. The original seven-marker reward with every gene clampable (the legacy policy) is kept in the code as an option and is used only in Fig. 4 and Supplementary Table 5. Rewards lie in [0, 1].

### Additive and Bliss nulls

For a set of components *d* with marker-level probabilities *p*_*d,i*_ of the desired state, untreated probabilities *u*_*i*_ and set score *s* (the mean over markers), the additive expectation is 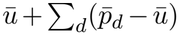 with bars denoting means over markers. The Bliss expectation is computed per marker from fractional effects *e*_*d,i*_ = (*p*_*d,i*_ *− u*_*i*_)*/*(1 *− u*_*i*_) as *u*_*i*_ + (1 *− u*_*i*_)[1 *−* Π _*d*_(1 *− e*_*d,i*_)] and then averaged over markers. The excess is the observed score minus the expectation. These are model-based nulls on a fraction-of-markers reward.

### Certified search

In round *t* each active candidate receives *m* more simulations (*m* = 100 in the benchmark, 200 in the druggable search), so every active candidate has *n*_*t*_ = *mt* samples. With sample mean 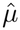 and unbiased sample variance 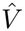, the radius is

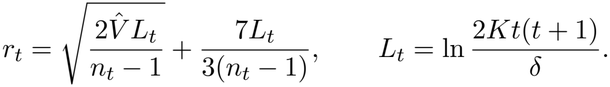

A candidate is eliminated when 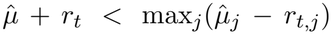. The search stops when LCB_*b*_ ≥ max_*j* ≠ *b*_ UCB_*j*_ *− ε* for the empirical best *b*, or when one candidate remains. Runs that reach 40,000 samples per candidate (60,000 in the druggable search) are flagged as uncertified.

#### Theorem 1

(informal; Supplementary Note 1). *With these constants, with probability at least* 1*−*2*δ all confidence intervals hold at once, an optimal candidate is never eliminated, and the returned candidate b satisfies µ*_*b*_ ≥ *µ*^*⋆*^ *− ε*.

The factor of two appears because *L*_*t*_ is the one-sided Maurer–Pontil constant. Replacing 2 by 4 inside the logarithm gives 1 *− δ*, and in 20 paired runs on the druggable family it cost 5.0% more simulations. The proof extends to any sampling rule when the union bound is taken over (candidate, number of batches) and each candidate has its own reward stream (Theorem 2), and Supplementary Note 1 gives an explicit-constant sample-complexity bound for successive elimination (Theorem 3, which also uses the sample-variance bound of [12]) that we checked numerically against all replayed runs.

### Baselines and ground truth

Each baseline received as many simulations as the matching certified run used. Uniform allocation splits them equally. Fixed-budget successive halving follows Karnin and colleagues [10]. Greedy forward selection adds the node with the best estimated score, up to the maximum set size, and returns the best set it visited. LUCB [8] samples the empirical best and the challenger with the highest upper bound, and UGapE [9] samples the one of its two leading candidates with the wider interval; both use the same empirical-Bernstein radius and cap (see below). For ground truth we screened all candidates with 2,000 simulations and re-estimated the top 30 with 60,000. The best refined estimate defines the optimum, and a returned set counts as correct if its refined score is within *ε* of it. To bound the chance that a set outside the top 30 was better, we used a union bound with the worst-case standard deviation 0.5 of a [0, 1] reward.

### Method comparison and ablations

Six families were replayed from simulated pools: the 92 druggable AML sets (100,000 simulations each), the 833 sets of up to three of the 17 clampable AML genes (40,000 each), the 15 of these with the highest pooled mean, and all candidate sets of the Grieco, Calzone and Remy networks under their pre-registered objectives (40,000 each). Each candidate’s rewards take the *q* + 1 values 0, 1*/q*, … , 1 for *q* reward markers, so a pool is a table of counts, and a draw of *m* rewards is a multinomial sample from the pooled distribution. This gives independent draws from a known distribution, whose mean is the exact ground truth. All procedures use *δ* = 0.05, the same batch size *m* (200 for the druggable family, 100 otherwise) and cap (100,000 and 40,000 samples per candidate) and the radius of the next paragraph, evaluated with

*L* = ln (2*Kj*(*j* + 1)*/δ*) where *j* is the number of batches a candidate has received. This union bound over (candidate, *j*) is valid for any sampling rule when each candidate has its own reward stream (the proof of Lemma 2 applies to the first *mj* draws of each candidate). Variants: *no elimination* keeps all candidates active; *Hoeffding* replaces the radius by 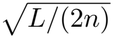 *doubling checkpoints* evaluates the stopping rule at *n* = *m*, 2*m*, 4*m*, … with *j* the checkpoint index and stops when the next checkpoint would exceed the cap. LUCB samples the empirical best and the challenger with the highest upper bound each round and stops under the certified-search rule; UGapE recommends the candidate *J* minimising *B*_*k*_ = max_*i* ≠ *k*_ UCB_*i*_ *−* LCB_*k*_, stops when *B*_*J*_ ≤ *ε* and otherwise samples whichever of *J* and the strongest challenger has the wider interval. Runs counted violations if any evaluated interval excluded the true mean, and as correct if the returned set was within *ε* of the best true mean. Batch LUCB samples, in each round, the empirical best candidate and the *B −* 1 other candidates with the highest upper bounds (*B* = 4, 16, 128), *m* simulations each, stops under the LUCB rule and is evaluated with *W* = *B* workers (20 to 100 runs per setting). Matched-budget uniform allocation and successive halving received the budget of the paired certified run. For elapsed time with *W* workers, a round in which *A* batches of *b* simulations are run costs ⌈*A/W* ⌉ *b* simulation-times; certified search runs one batch per active candidate per round, LUCB two and UGapE one, after an initial round of *K* batches. Settings and runs: Table 2. For the controlled test, five sub-families per row were drawn from the 833 sets, always including the *j* best, with six runs each (*ε* = 0.02).

### Calibration ensemble and structural sensitivity

The calibrations are all distinct KGBN calibrations with the same 24 genes (duplicates removed by file hash). Their dependence was summarised by the pairwise Spearman correlation of rule-probability vectors and of the scores of the ten five-drug candidates. Structural perturbations act on the rules of the reference network: removing the MAPK1 token from the STAT5A rules, swapping the probabilities of the two main STAT5A rules, and replacing a randomly chosen regulator token in a randomly chosen rule by a random gene (one or three such rewirings; 60 networks each). Each perturbed network was evaluated with 4,000–8,000 simulations per set.

### Other AML analyses

The size-matched null enumerates all one-, two- and three-gene sets over the 17 clampable genes (1,500 simulations each). Ranking stability uses 200 independent batches. Genotype strata hold mutation status fixed at every step with drug clamps taking precedence. Rule-probability perturbation draws each gene’s probability vector from a Dirichlet distribution with concentration *κ* times the fitted vector. The scores of the five-drug candidates come from Hoeffding-sized runs with (*ε, δ*) = (0.01, 0.01), i.e. *n* = 26,492.

### Extended analyses

*Audit:* true scores of the 92 druggable sets from 60,000 simulations each; 60 runs with seeds 5000–5059; a violation is any round at which an active set’s empirical mean differs from its true score by more than its radius. The harder audit used the 15 AML gene sets with the highest screening scores, *ε* = 0.02 and 15 runs. *Sensitivity:* six (*ε, δ*) settings, six runs each (cap 100,000 samples per set). *Bound ablation:* the empirical-Bernstein radius against 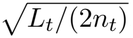, three runs each at *ε* = 0.02 (cap 40,000). *Scaling:* sub-families drawn without replacement from the AML gene sets, three per size (the controlled test above separates size from near-ties); near-ties counted in a 1,500-simulation screen of the whole family. *Noise:* Calzone with maximum set size two (*K* = 351), three runs per noise level, ground truth as in the benchmark. *Genotype:* FLT3 and NPM1 held on or off; each stratum searched with *ε* = 0.01 and ranked from a 20,000-simulation reference per set. *Marker decomposition:* 40,000–60,000 simulations per component, pair and the triplet, with marker-level probabilities taken from the final state.

### Decoy maps and loss of function

For each decoy map, every one of the eight drug units received a random set of clampable genes of the same size, drawn without replacement (so unit overlaps such as the shared FLT3 target of the real map are not preserved), and the 92-set search was repeated with 1,000 simulations per set. The real map was evaluated 30 times with the same procedure to show its noise, and we report the mean. One-sided *p*-values are (1 + #*{*decoys ≥ real*}*)*/*(1 + #decoys), computed with the mean of the replicate evaluations of the real map, and are adjusted over the four reported statistics by the Holm method. The decoy analysis was not pre-specified. For loss of function, one gene was held off throughout (also when untreated) with 1,500 simulations per set; the reward genes and the mutation nodes were not removed. Stage `G_decoy` of the code produces these results.

### Out-of-sample check

We used only evaluations shipped with KGBN. Leave-one-drug-out (5 folds, test *n* = 8 genotypes) and leave-one-profile-out (8 folds, test *n* = 5 drugs) models were scored on their held-out points; the Waves 3+4 and Waves 1+2 fits were scored on the other cohort and on the FPMTB cohort. Agreement is Spearman’s *ρ* between the network output and measured AUC. For FPMTB, which scores sensitivity (higher is more sensitive) while AUC and the network output rise with resistance, the sign was reversed. Baselines: for a held-out drug, the mean AUC of the other four drugs at the same genotype; for a held-out genotype, the mean AUC of the same drug at the other genotypes; across cohorts, the measured training-cohort values (persistence), which a perfect fit to the training cohort would reproduce. Permutation *p*-values are exact for *n* ≤ 8 and use 20,000 permutations otherwise; intervals are bootstrap percentile intervals (4,000 resamples of points). Points from different folds are not independent, so the pooled intervals are approximate. Stage F validation of the code produces these results.

### Pre-specification and deviations

We wrote the benchmark protocol before running it. There were two deviations from it. Remy tumorigenesis was stopped after three repetitions because of run time, and the T-LGL network was dropped because its untreated score (0.925) left no headroom. Two later changes were made after review: the AML network was re-analysed under the strict policy described above, and the repetitions for the AML, Calzone and Grieco networks were increased to eight.

## Data and code availability

All inputs are public: the KGBN repository (https://github.com/IlyaLab/KGBN) and the PyBoolNet repository (https://github.com/hklarner/pyboolnet). A single Python file, aml certified search.py, generates the tables and figures in this paper. No notebooks are included, because earlier notebooks contain the superseded policy. Result tables and the pre-specified protocol are supplementary files. The code is available at https://github.com/wasimrazaa135-ops/aml-certified-search and is archived on Zenodo at https://doi.org/10.5281/zenodo.24450270.

## Author contributions and competing interests

W.R.: conceptualization, methodology, software, formal analysis, investigation, writing – original draft. S.M.: formal analysis (theory and proofs), validation, writing – review and editing. C.E.: investigation, visualization, writing – review and editing. F.A.B.d.S.: supervision, resources, writing – review and editing. All authors approved the final manuscript. F.A.B.d.S. co-authored the earlier pipeline [4] on which this work builds. The authors declare no other competing interests.

## Declaration of Generative AI and AI-Assisted Technologies

The authors declare that no generative AI or AI-assisted technologies were used in the analysis or writing of this manuscript.

## Supplementary information

### Supplementary Note 1: correctness and cost of certified search

#### Setting

A finite family *A* with |*A*| = *K*; i.i.d. rewards *X* ∈ [0, 1] for each candidate, with mean *µ*_*a*_; *µ*^*⋆*^ = max_*a*_ *µ*_*a*_.

##### Lemma 1

**(empirical Bernstein [12])**. For i.i.d. *X*_*i*_ ∈ [0, 1] and *n* ≥ 2, with probability at least 1 *− δ*^*′*^,

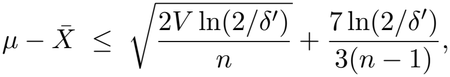

and the symmetric bound holds for 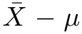, where *V* is the unbiased sample variance. Our radius divides the variance term by *n −* 1 rather than *n*, which is slightly larger and so conservative.

#### Reward-stream model

Each candidate *a* has an i.i.d. sequence *X*_*a*,1_, *X*_*a*,2_, … , independent across candidates, and a procedure that has drawn *n* samples from *a* has seen the first *n* entries. Whatever the sampling rule, the mean and variance of the first *n* entries are those of a fixed-size sample.

##### Lemma 2.

Let *r*_*j*_(*a*) denote the radius of Lemma 1 with *δ*^*′*^ = *δ*_*a,j*_ = *δ/*(*Kj*(*j* + 1)) applied to the first *mj* samples of candidate *a*, that is *L*_*j*_ = ln (2*Kj*(*j* + 1)*/δ*), and let *G* be the event that 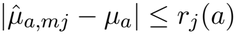 for every candidate *a* and every *j* ≥ 1. Then Pr(*G*^*c*^) ≤ 2*δ*.

*Proof*. For a fixed pair (*a, j*) each tail fails with probability at most *δ*_*a,j*_, because the first *mj* entries form a fixed-size i.i.d. sample. Summing, ∑_*j*_ ∑_*a*_ 2*δ*_*a,j*_ = 2*K* ∑_*j*_ *δ/*(*Kj*(*j* + 1)) = 2*δ*.

Successive elimination has the same *j* = *t* for every active candidate, so its *L*_*t*_ is the *L*_*j*_ above with *j* = *t*. LUCB, UGapE and batch LUCB use *j* = *n*_*a*_*/m* for candidate *a*.

##### Theorem 1

**(successive elimination)**. On *G* the procedure never eliminates an optimal candidate, and any returned candidate *b* satisfies *µ*_*b*_ ≥ *µ*^*⋆*^ *− ε*.

*Proof*. To eliminate an optimal *a*^*⋆*^ the procedure would need 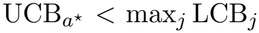. On *G*, though, 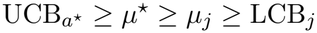 for every *j*. At a stopping time with LCB_*b*_ ≥ UCB_*j*_ *− ε* for all active *j* ≠ *b*, and *a*^*⋆*^ active, on *G* we have 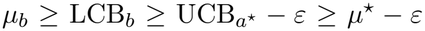. If a single candidate remains it is optimal.

##### Theorem 2

**(index methods, including batch LUCB)**. Under any sampling rule, if LUCB (or batch LUCB) stops with LCB_*h*_ ≥ max_*l* ≠ *h*_ UCB_*l*_ *− ε* for the empirical best *h*, or UGapE stops with max_*i* ≠ *J*_ UCB_*i*_ *−* LCB_*J*_ ≤ *ε*, then on *G* the returned candidate satisfies *µ*_*h*_ ≥ *µ*^*⋆*^ *− ε* (respectively *µ*_*J*_ ≥ *µ*^*⋆*^ *− ε*). Hence it is *ε*-optimal with probability at least 1 *−* 2*δ*.

*Proof*. On *G*, 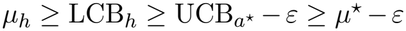 when *h* ≠ *a*^*⋆*^, and trivially when *h* = *a*^*⋆*^. The UGapE case is the same with *J*. Only *G* is used, so the sampling rule is arbitrary.

#### Cost of the stricter constant

Using *L*_*t*_ = ln(4*Kt*(*t* + 1)*/δ*) makes each two-sided interval fail with probability at most 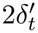 with 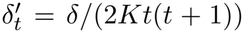, giving Pr(*G*^*c*^) ≤ *δ*. The radius grows by a factor of about 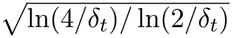, roughly 1.03 at the scales used here, and because cost scales with the square of the radius the expected increase in simulations is about 5%. In 20 paired runs on the 92-set family the measured increase was 5.0%.

##### Theorem 3

**(sample complexity of successive elimination, explicit constants)**. Let 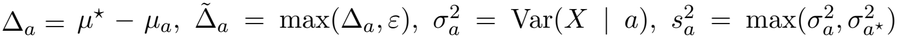 and *m* ≥ 2. Let 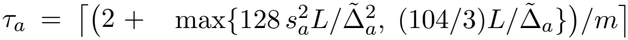 and 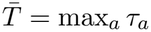, with 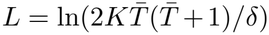 (a fixed point of 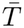). With probability at least 1 *−* 3*δ*, successive elimination stops after at most 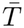 rounds and draws at most *mτ*_*a*_ samples from every candidate with Δ_*a*_ *> ε* and at most 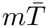 samples from every candidate with Δ_*a*_ ≤ *ε*, so

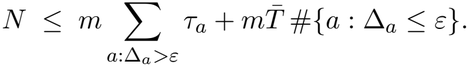

*Proof*. Let *V* be the event that 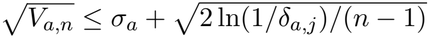 for all (*a, j*), *n* = *mj*, where *V*_*a,n*_ is the unbiased sample variance; by Theorem 10 of Maurer and Pontil, Pr(*V* ^*c*^) ≤ ∑_*a,j*_ *δ*_*a,j*_ = *δ*, so Pr((*G ∩ V*)^*c*^) ≤ 3*δ*. On *V* , with *l* = ln(1*/δ*_*a,j*_) ≤ *L*_*j*_, the radius satisfies 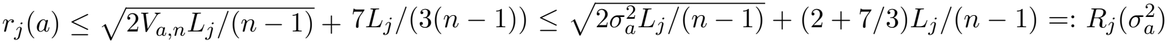. (i) A candidate *j* with Δ_*j*_ *> ε* is eliminated as soon as 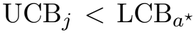. On *G*, UCB_*j*_ ≤ *µ*_*j*_ + 2*r*_*j*_(*j*) and 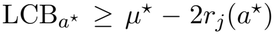, so it suffices that *r*_*j*_(*j*) *<* Δ_*j*_*/*4 and *r*_*j*_(*a*^*⋆*^) *<* Δ_*j*_*/*4. Because *R* increases in *σ*^2^ and the two terms of *R*_*j*_ *< x* are each below *x/*2 when *n −* 1 *>* 8*σ*^2^*L/x*^2^ and *n −* 1 *>* (26*/*3)*L/x*, taking *x* = Δ_*j*_*/*4 gives the conditions 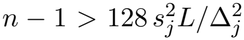 and *n −* 1 *>* (104*/*3)*L/*Δ_*j*_, which hold at round *τ*_*j*_ (and *L*_*j*_ ≤ *L* for all rounds up to 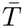). (ii) At a round at which every candidate with Δ *> ε* has been eliminated and every remaining candidate has 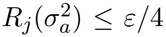, the stopping rule holds: for the empirical best *b*, 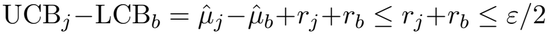. That requires *n−*1 ≥ 32*σ*^2^*L/ε*^2^ and *n−*1 ≥ (52*/*3)*L/ε*, which the definition of 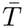 implies.

Uniform allocation with the same guarantee needs *K*⌈ln(2*K/δ*)*/*(2*ε*^2^)⌉. The bound has the usual form 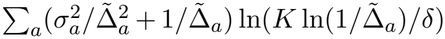, with the variance of the optimal candidate entering every term, and it is not smaller than the uniform design for large families with many near-ties (the Remy network). We evaluated it on all replayed families (Table 4): it held in 11/11 settings and every run, and it was 7 to 31 times larger than the simulations used. The radius inequality 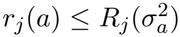 was exceeded in 0 of 80,000 checks on random streams from 100 AML candidates. These checks do not replace the proof. We have not proved a sample-complexity bound for batch LUCB, UGapE or LUCB under this confidence sequence.

**Table 4:** Numerical check of the explicit sample-complexity bound (Theorem 3). The bound is evaluated from the true means and variances of each replayed family and compared with the simulations used by successive elimination in the runs of Table 20.

| Family | ε | Runs | Mean observed | Largest observed | Theorem 3 bound | Bound / mean | Holds in every run |
|---|---|---|---|---|---|---|---|
| AML 3-gene sets (K=833) | 0.02 | 50 | 597,664 | 661,600 | 5,366,600 | 9× | yes |
| AML 3-gene sets (K=833) | 0.05 | 50 | 546,998 | 609,300 | 3,959,400 | 7× | yes |
| AML druggable sets (K=92) | 0.01 | 100 | 79,762 | 91,800 | 2,489,000 | 31× | yes |
| AML druggable sets (K=92) | 0.02 | 100 | 71,564 | 85,200 | 1,182,200 | 17× | yes |
| AML druggable sets (K=92) | 0.05 | 100 | 62,586 | 73,000 | 540,000 | 9× | yes |
| Calzone cell fate (K=2,951) | 0.02 | 30 | 3,321,077 | 3,551,600 | 40,625,300 | 12× | yes |
| Calzone cell fate (K=2,951) | 0.05 | 30 | 2,560,553 | 2,722,600 | 25,729,000 | 10× | yes |
| Grieco MAPK (K=1,275) | 0.02 | 30 | 1,388,680 | 1,461,300 | 18,484,500 | 13× | yes |
| Grieco MAPK (K=1,275) | 0.05 | 30 | 1,209,453 | 1,291,500 | 13,071,900 | 11× | yes |
| Remy tumorigenesis (K=5,488) | 0.02 | 20 | 13,161,835 | 14,268,400 | 318,844,700 | 24× | yes |
| Remy tumorigenesis (K=5,488) | 0.05 | 20 | 7,840,070 | 8,568,200 | 151,682,300 | 19× | yes |

### Supplementary Note 2: the five-drug analysis under the strict policy

The untreated score of the AML network is 44.7%. The best pair (midostaurin + ibrutinib) scores 53.4%; its strict, all-markers-correct success rate is only 3.3%, which is why we use a continuous score. Midostaurin + entospletinib is sub-additive by -1.3 points. The effect of the clamping policy is in Table 5 and Fig. 4. The pair’s excess under the additive and Bliss nulls is in Table 6.

**Table 5:** Effect of the clamping policy. Gain over the untreated score (percentage points) for the same simulations scored with the original seven-marker reward (which includes the drug targets BCL2 and AKT1) and the five-marker reward used in the main analysis. No gene rule uses BCL2 as a regulator, so under the strict reward venetoclax cannot change any output.

| Candidate | Legacy 7-marker gain (pp) | Strict 5-marker gain (pp) |
|---|---|---|
| Venetoclax (BCL2) | +10.9 | +0.0 |
| Ibrutinib (BTK) | +2.2 | +3.1 |
| Entospletinib (SYK) | +10.5 | +4.0 |
| Selumetinib (MAP2K1/2) | +1.6 | +1.5 |
| Midostaurin (FLT3+KIT) | +8.7 | +1.5 |
| AKT-inhibitor class (AKT1) | +6.4 | -1.6 |
| ERK-inhibitor class (MAPK1) | +7.9 | +7.0 |
| Midostaurin + Venetoclax | +19.1 | +1.5 |
| Midostaurin + Ibrutinib | +13.9 | +8.8 |
| Ibrutinib + Venetoclax | +13.1 | +3.1 |
| BCL2 + MYC + SYK (best 3-gene set under legacy policy) | +37.7 | +27.6 |

**Table 6:** **Five-drug pairs under the strict reward**, with linear-additive and Bliss expectations.

| Pair | Observed (%) | Additive (%) | Bliss (%) | Excess additive (pp) | Excess Bliss (pp) |
|---|---|---|---|---|---|
| Midostaurin + Venetoclax | 46.2 | 46.2 | 46.2 | +0.00 | +0.00 |
| Midostaurin + Ibrutinib | 53.4 | 49.2 | 48.9 | +4.21 | +4.54 |
| Midostaurin + Entospletinib | 48.8 | 50.1 | 49.7 | -1.34 | -0.89 |
| Midostaurin + Selumetinib | 48.0 | 47.7 | 47.6 | +0.34 | +0.48 |
| Ibrutinib + Venetoclax | 47.7 | 47.7 | 47.7 | +0.00 | +0.00 |

### Supplementary Note 3: patient-level uncertainty

The public KGBN repository contains per-patient BeatAML venetoclax AUC values (352 samples) with mutation profiles. The values are not normally distributed within groups (Shapiro–Wilk *p <* 0.05 in four of eight profiles). We compared a bootstrap confidence interval for the mean, a Gaussian prediction interval, a residual-bootstrap interval, split conformal prediction [21, 22], conformalised quantile regression [23], Mondrian (profile-conditional) conformal prediction and covariate-shift-weighted conformal prediction [24] under a random split, leave-one-mutation-profile-out, and a real cohort shift between BeatAML waves (Fig. SS1). Intervals targeted 90% coverage; the linear model used three mutation indicators; the designs used 150, 100 and 100 repetitions.

A bootstrap interval for the mean covered 9.2% of held-out patients in the leave-one-profile-out design (10.3% under a random split). Proper prediction intervals do not fail this way, and conformal prediction did not do better than they did: coverage in the leave-one-profile-out design was 92.5% for a Gaussian prediction interval, 90.2% for split conformal (SD 6.9 points across repeated splits), 86.7% for a residual-bootstrap interval and 82.2% for conformalised quantile regression (SD 19.2 points). Under the real cohort shift, split conformal covered 87.8% and 92.2%. Marginal coverage hid differences between subgroups: under a random split the lowest profile-conditional coverage was 86.6% for split conformal and 89.3% for Mondrian conformal. All valid intervals were wide, about 253 AUC units against 25 for the bootstrap interval of the mean, because three mutation indicators explain little of the variation between individuals. The pooled correlation between the calibrated network and observed BeatAML response is 0.879 over 40 drug-by-genotype points; within a single drug across eight genotypes it ranges from 0.29 (midostaurin) to 0.94 (venetoclax).

**Figure S1:**
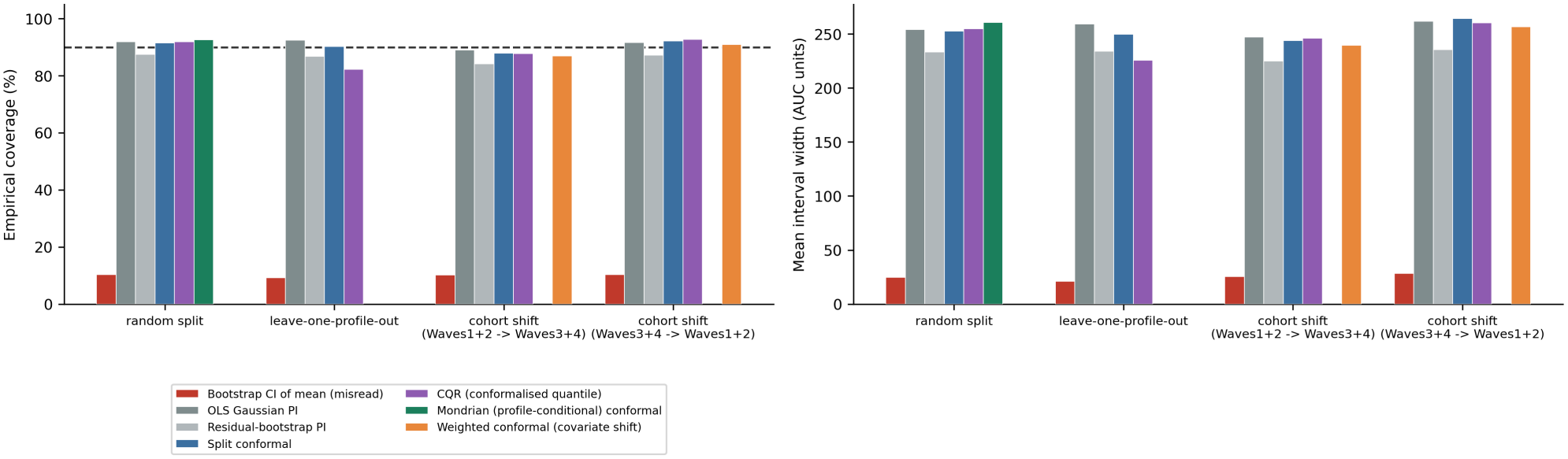
Real per-patient BeatAML venetoclax AUC: prediction intervals under four designs. Left, empirical coverage; right, mean interval width. The dashed line is the 90% target. The bootstrap interval of the mean is shown only to illustrate how it is misread.

### Supplementary Note 4: a topological ranking criterion did not hold up

We initially ranked candidates by the total persistence of one-dimensional homology (the Topological Bottleneck Score, TBS) in the cloud of transient states visited during simulated inhibition, and read lower scores as better. We tested it against exact or Monte-Carlo ground truth over large candidate universes (66, 136 and 253 sets). TBS correlated *positively* with outcome (Spearman *ρ* = 0.64, 0.51 and 0.26 in the pilot, Palma and AML networks), the opposite of our interpretation. After controlling for trajectory length or cloud size the partial correlations were 0.06, 0.27 and -0.08. On the pilot network, mean transient length predicted exact convergence better (*ρ* = 0.84), and a static structural heuristic predicted the exact functional score far better than TBS (*ρ* = 0.91 against 0.37). TBS also depended on the seed set: the test–retest correlation was 0.69 at 40 seeds and 0.93 at 160, and the TBS-minimising candidate was the same in only 20% of seed draws. The AML-network analysis (253 sets) uses the strict reward. We do not recommend TBS as a ranking criterion; we report this negative result so that others can avoid the same approach.

**Figure S2:**
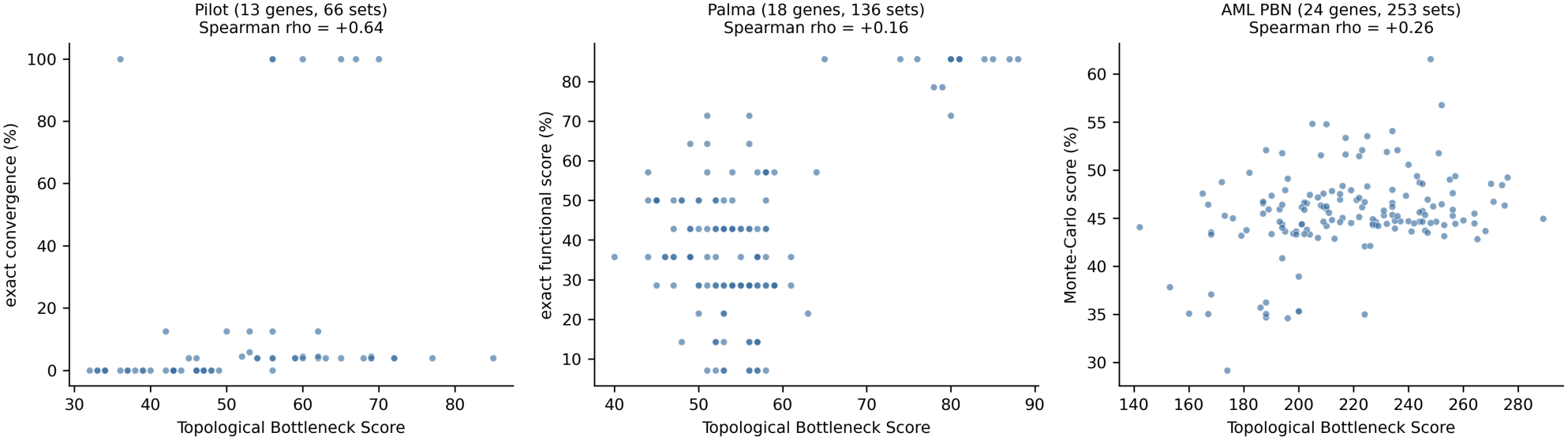
TBS against ground truth in three networks. Each point is one candidate intervention.

### Supplementary Note 5: complete benchmark and application tables

Table 7 gives the reward definitions fixed before the benchmark. Table 8 lists every benchmark run, Table 9 the ground-truth margin and Table 10 the fifteen highest-scoring druggable combinations. Table 11 gives the full 92-set search inside every calibration, Table 12 the structural sensitivity analysis and Table 18 the marker-level ablation.

**Table 7:** Reward definition and family size for each network (pre-specified). Reward is the fraction of the listed output nodes in the desired state at the final step.

| Network | Desired ON | Desired OFF | Max set size |
|---|---|---|---|
| Calzone cell fate | Apoptosis | Survival | 3 |
| Remy tumorigenesis | Apoptosis_high, Growth_arrest | Proliferation | 3 |
| Grieco MAPK | Apoptosis, Growth_Arrest | Proliferation | 2 |
| Zhang T-LGL leukemia (excluded) | Apoptosis | Proliferation | 2 |
| AML PBN (KGBN) | TP53, CDKN2A | BCL2, MYC, CCND1, AKT1, STAT5A | 3 |

**Table 8:** Every benchmark run. Simulations and reduction relative to the guaranteed-uniform design; the last three columns say whether each budget-matched baseline returned an *ε*-optimal set.

| Network | Run | Simulations | Reduction (×) | Returned set | Certified correct | Uniform | Halving | Greedy |
|---|---|---|---|---|---|---|---|---|
| AML PBN | 1 | 612,000 | 17.7 | BTK+MAPK1+SYK | yes | yes | yes | yes |
| AML PBN | 2 | 610,800 | 17.8 | BTK+MAPK1+SYK | yes | yes | yes | yes |
| AML PBN | 3 | 628,600 | 17.3 | BTK+FLT3+MAPK1 | yes | yes | yes | yes |
| AML PBN | 4 | 617,300 | 17.6 | BTK+MAPK1+SYK | yes | yes | yes | yes |
| AML PBN | 5 | 570,900 | 19.0 | BTK+MAPK1+SYK | yes | yes | yes | yes |
| AML PBN | 6 | 573,400 | 18.9 | BTK+MAPK1+SYK | yes | yes | yes | yes |
| AML PBN | 7 | 614,900 | 17.6 | BTK+MAPK1+SYK | yes | yes | yes | yes |
| AML PBN | 8 | 601,200 | 18.0 | BTK+FLT3+MAPK1 | yes | yes | yes | yes |
| Calzone | 1 | 3,390,300 | 12.7 | MPT+NFkB+XIAP | yes | no | yes | no |
| Calzone | 2 | 3,067,100 | 14.0 | MPT+NFkB+XIAP | yes | yes | yes | no |
| Calzone | 3 | 3,243,700 | 13.3 | MPT+NFkB+XIAP | yes | yes | yes | no |
| Calzone | 4 | 3,211,500 | 13.4 | MPT+NFkB+XIAP | yes | yes | yes | no |
| Calzone | 5 | 3,052,300 | 14.1 | MPT+NFkB+XIAP | yes | yes | yes | no |
| Grieco | 1 | 1,431,200 | 12.1 | AKT+DUSP1 | yes | yes | yes | yes |
| Grieco | 2 | 1,316,400 | 13.1 | AKT+DUSP1 | yes | yes | yes | yes |
| Grieco | 3 | 1,408,700 | 12.3 | AKT+DUSP1 | yes | yes | yes | yes |
| Grieco | 4 | 1,405,900 | 12.3 | AKT+DUSP1 | yes | yes | yes | yes |
| Grieco | 5 | 1,388,200 | 12.4 | AKT+DUSP1 | yes | yes | yes | yes |
| Remy | 1 | 13,166,200 | 6.4 | CyclinA+CyclinD1+CyclinE1 | yes | yes | yes | yes |
| Remy | 2 | 12,986,800 | 6.5 | CyclinA+CyclinD1+CyclinE1 | yes | yes | yes | yes |
| Remy | 3 | 13,124,800 | 6.4 | CyclinA+CyclinD1+CyclinE1 | yes | yes | yes | yes |

**Table 9:** Ground-truth margin. A set outside the re-estimated top 30 would have to beat the best refined estimate and still fall below the 30th screening score; the union bound uses the worst-case standard deviation 0.5 of a [0,1] reward.

| Network | *K* | Best refined | 30th screening score | Margin | Union bound on missing a better set |
|---|---|---|---|---|---|
| AML PBN (KGBN) | 833 | 0.684 | 0.558 | 0.126 | 1.1e-26 |
| Calzone cell fate | 2951 | 0.809 | 0.776 | 0.034 | 1 |
| Grieco MAPK | 1275 | 0.877 | 0.815 | 0.063 | 1.4e-05 |

**Table 10:** Fifteen highest-scoring druggable combinations. Score is the model score from a 30,000-simulation reference. Additive: untreated score plus summed single-drug gains. Bliss: marker-level Bliss independence of fractional effects, averaged over markers. FLT3i: gilteritinib-class FLT3 inhibitor; ERKi, AKTi: inhibitor classes mapped to MAPK1 and AKT1.

| Rank | Combination | Score | Additive | Excess (additive) | Bliss | Excess (Bliss) |
|---|---|---|---|---|---|---|
| 1 | Ibrutinib + Midostaurin + ERKi | 0.710 | 0.560 | +0.149 | 0.551 | +0.158 |
| 2 | Ibrutinib + Entospletinib + ERKi | 0.683 | 0.586 | +0.097 | 0.569 | +0.115 |
| 3 | Ibrutinib + FLT3i + ERKi | 0.683 | 0.561 | +0.123 | 0.551 | +0.132 |
| 4 | Entospletinib + Midostaurin + ERKi | 0.642 | 0.568 | +0.074 | 0.557 | +0.085 |
| 5 | Entospletinib + AKTi + ERKi | 0.610 | 0.542 | +0.068 | 0.540 | +0.070 |
| 6 | Venetoclax + Entospletinib + ERKi | 0.609 | 0.557 | +0.052 | 0.551 | +0.058 |
| 7 | Entospletinib + ERKi | 0.609 | 0.555 | +0.054 | 0.550 | +0.059 |
| 8 | Entospletinib + FLT3i + ERKi | 0.608 | 0.568 | +0.040 | 0.557 | +0.051 |
| 9 | Midostaurin + AKTi + ERKi | 0.602 | 0.516 | +0.086 | 0.517 | +0.086 |
| 10 | Midostaurin + ERKi | 0.599 | 0.529 | +0.069 | 0.528 | +0.071 |
| 11 | Venetoclax + Midostaurin + ERKi | 0.598 | 0.531 | +0.067 | 0.529 | +0.069 |
| 12 | Midostaurin + FLT3i + ERKi | 0.598 | 0.542 | +0.056 | 0.537 | +0.061 |
| 13 | Ibrutinib + Selumetinib + Midostaurin | 0.590 | 0.505 | +0.085 | 0.498 | +0.092 |
| 14 | Ibrutinib + Entospletinib + Selumetinib | 0.586 | 0.531 | +0.055 | 0.517 | +0.070 |
| 15 | Ibrutinib + Selumetinib + FLT3i | 0.585 | 0.506 | +0.080 | 0.498 | +0.087 |

**Table 11:** Certified search inside every distinct calibration. The full 92-set druggable search was repeated in each calibration shipped with the KGBN toolkit (duplicates removed by file hash). The triplet is ibrutinib + midostaurin + ERK-inhibitor class; its rank is by a 3,000-simulation reference. These calibrations share rule sets and part of their training data and are not statistically independent. LODO: leave-one-drug-out; LOCO: leave-one-cohort-out.

| Calibration | Triplet rank | Triplet certified best | Excess (additive) | Excess (Bliss) | Kendall τ of drug ranking |
|---|---|---|---|---|---|
| Waves3+4_AND+OR+IW | 1 | yes | +0.145 | +0.155 | 1.00 |
| Waves1+2_AND+OR+IW/FPMTB_sDSS_5drugs | 1 | yes | +0.161 | +0.195 | 0.64 |
| LOCO_fold_0 | 1 | yes | +0.188 | +0.206 | 0.56 |
| LOCO_fold_1 | 1 | yes | +0.158 | +0.178 | 0.56 |
| LOCO_fold_2 | 1 | yes | +0.182 | +0.199 | 0.56 |
| LOCO_fold_3 | 1 | yes | +0.211 | +0.227 | 0.69 |
| LOCO_fold_4 | 1 | yes | +0.164 | +0.175 | 0.91 |
| LOCO_fold_5 | 1 | yes | +0.342 | +0.356 | 0.73 |
| LOCO_fold_6 | 1 | yes | +0.225 | +0.247 | 0.64 |
| LOCO_fold_7 | 1 | yes | +0.101 | +0.137 | 0.56 |
| LODO_fold_0 | 1 | yes | +0.083 | +0.107 | 0.64 |
| LODO_fold_1 | 1 | yes | +0.152 | +0.166 | 0.60 |
| LODO_fold_2 | 1 | yes | +0.194 | +0.210 | 0.73 |
| LODO_fold_3 | 1 | yes | +0.173 | +0.194 | 0.60 |
| LODO_fold_4 | 1 | yes | +0.247 | +0.272 | 0.73 |

**Table 12:** Structural sensitivity of the triplet. Excess over additivity of ibrutinib + midostaurin + ERK-inhibitor class when the MAPK1 input to STAT5A is removed, when the two main STAT5A rule probabilities are swapped, and under random rewiring of one or three regulators. The last column counts networks in which the triplet scores highest among the four triplets that can be formed from ibrutinib, midostaurin, entospletinib and the ERK-inhibitor class.

| Scenario | Networks | Mean excess (additive) | Minimum | Mean excess (Bliss) | Excess > 0.02 | Triplet best of four triplets |
|---|---|---|---|---|---|---|
| MAPK1 input to STAT5A removed | 1 | +0.143 | +0.143 | +0.150 | 100% | 100% |
| STAT5A OR-rule and AND-rule probabilities swapped | 1 | +0.089 | +0.089 | +0.107 | 100% | 100% |
| random rewiring of 1 regulator(s) | 60 | +0.141 | +0.077 | +0.150 | 100% | 100% |
| random rewiring of 3 regulator(s) | 60 | +0.132 | +0.056 | +0.143 | 100% | 98% |
| reference network | 1 | +0.152 | +0.152 | +0.161 | 100% | 100% |

### Supplementary Note 6: extended-analysis tables

Tables 13–17 and Table 19 give the numbers behind Fig. 3 and Fig. 9. Tables 20–22 and 23 give the method comparison, the elapsed-time results, the batch-size ablation and the controlled gap-structure test. (An earlier four-run comparison with LUCB, which indexed the radius by round and not by candidate, is superseded by Table 20.)

**Table 13:** Cost against accuracy and confidence targets. (92 druggable combinations).

| ε | δ | Runs | Certified | ε-optimal | Mean simulations | Uniform bound | Reduction (×) |
|---|---|---|---|---|---|---|---|
| 0.005 | 0.05 | 6 | 6 | 6 | 83,767 | 15,107,688 | 180.8 |
| 0.01 | 0.01 | 6 | 6 | 6 | 87,100 | 4,517,292 | 52.0 |
| 0.01 | 0.05 | 6 | 6 | 6 | 77,533 | 3,776,968 | 48.8 |
| 0.01 | 0.2 | 6 | 6 | 6 | 68,967 | 3,139,224 | 45.6 |
| 0.02 | 0.05 | 6 | 6 | 6 | 70,033 | 944,288 | 13.5 |
| 0.05 | 0.05 | 6 | 6 | 6 | 61,700 | 151,156 | 2.5 |

**Table 14:** Scaling with family size. Random sub-families of the 1,793 AML gene sets; near-ties are sets within 0.04 of the best in a 1,500-simulation screen.

| *K* | Certified | Near-ties (mean) | Reduction mean (×) | Range |
|---|---|---|---|---|
| 50 | 3/3 | 3.7 | 6.8 | 4.0–8.6 |
| 100 | 3/3 | 1.0 | 18.0 | 15.0–20.9 |
| 250 | 3/3 | 3.7 | 13.3 | 6.4–18.7 |
| 500 | 3/3 | 3.7 | 16.5 | 11.3–20.9 |
| 800 | 3/3 | 2.7 | 18.8 | 18.1–19.3 |

**Table 15:** Noise sensitivity. (Calzone network, sets of up to two nodes).

| Noise | *K* | Untreated | Best | Certified | *ε*-optimal | Reduction (×) |
|---|---|---|---|---|---|---|
| 0.005 | 351 | 0.604 | 0.794 | 3/3 | 3/3 | 6.8 |
| 0.02 | 351 | 0.608 | 0.788 | 3/3 | 3/3 | 10.3 |
| 0.05 | 351 | 0.577 | 0.724 | 3/3 | 3/3 | 8.6 |

**Table 16:**
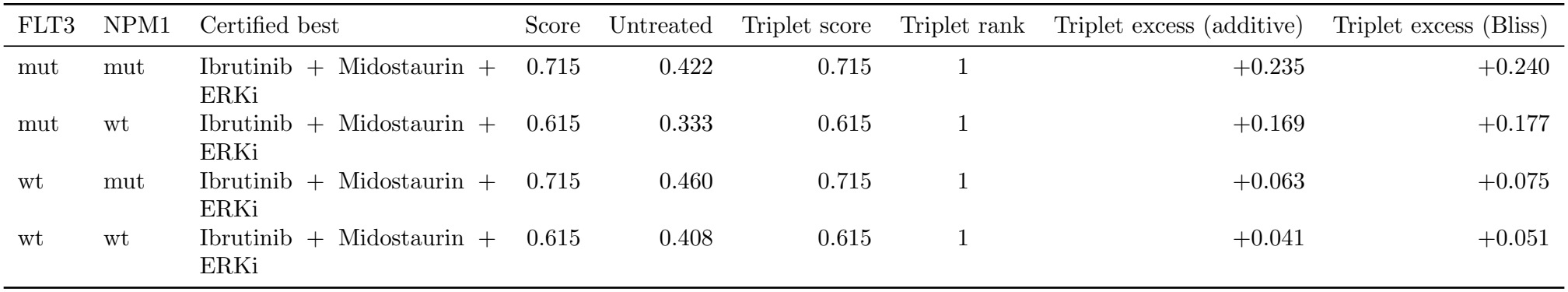
Genotype-conditional certified search. Mutation status held fixed; the triplet is ibrutinib + midostaurin + ERK-inhibitor class.

| FLT3 | NPM1 | Certified best | Score | Untreated | Triplet score | Triplet rank | Triplet excess (additive) | Triplet excess (Bliss) |
|---|---|---|---|---|---|---|---|---|
| mut | mut | Ibrutinib + Midostaurin + ERKi | 0.715 | 0.422 | 0.715 | 1 | +0.235 | +0.240 |
| mut | wt | Ibrutinib + Midostaurin + ERKi | 0.615 | 0.333 | 0.615 | 1 | +0.169 | +0.177 |
| wt | mut | Ibrutinib + Midostaurin + ERKi | 0.715 | 0.460 | 0.715 | 1 | +0.063 | +0.075 |
| wt | wt | Ibrutinib + Midostaurin + ERKi | 0.615 | 0.408 | 0.615 | 1 | +0.041 | +0.051 |

**Table 17:** Marker-level decomposition of the triplet. Independence expectation: *u* +(1 *− u*) [1 − Π_*d*_(1 *− g*_*d*_)] with *g*_*d*_ the single-component gain in marker probability.

| Marker (desired state) | Untreated | Independence | Observed triplet | Excess |
|---|---|---|---|---|
| TP53 on | 0.359 | 0.429 | 0.568 | +0.140 |
| CDKN2A on | 0.463 | 0.586 | 0.760 | +0.174 |
| MYC off | 0.438 | 0.522 | 0.650 | +0.128 |
| CCND1 off | 0.555 | 0.563 | 0.576 | +0.013 |
| STAT5A off | 0.418 | 0.656 | 1.000 | +0.344 |

**Table 18:** Marker-level ablation of the triplet. Marker probabilities of the desired state in the reference network and after removing the MAPK1 input, or both the BTK and MAPK1 inputs, from the STAT5A rules (40,000 simulations per set). The triplet is ibrutinib + midostaurin + ERK-inhibitor class.

| Network | Marker (desired state) | Untreated | Triplet | Additive | Excess (additive) | Bliss | Excess (Bliss) |
|---|---|---|---|---|---|---|---|
| reference network | TP53 on | 0.360 | 0.571 | 0.432 | +0.139 | 0.430 | +0.141 |
| reference network | CDKN2A on | 0.464 | 0.764 | 0.593 | +0.171 | 0.585 | +0.178 |
| reference network | MYC off | 0.435 | 0.648 | 0.521 | +0.127 | 0.521 | +0.127 |
| reference network | CCND1 off | 0.555 | 0.578 | 0.564 | +0.014 | 0.567 | +0.011 |
| reference network | STAT5A off | 0.421 | 1.000 | 0.698 | +0.302 | 0.658 | +0.342 |
| MAPK1 input to STAT5A removed | TP53 on | 0.366 | 0.571 | 0.438 | +0.133 | 0.436 | +0.135 |
| MAPK1 input to STAT5A removed | CDKN2A on | 0.471 | 0.764 | 0.602 | +0.161 | 0.593 | +0.171 |
| MAPK1 input to STAT5A removed | MYC off | 0.435 | 0.648 | 0.521 | +0.127 | 0.521 | +0.127 |
| MAPK1 input to STAT5A removed | CCND1 off | 0.547 | 0.578 | 0.555 | +0.023 | 0.562 | +0.015 |
| MAPK1 input to STAT5A removed | STAT5A off | 0.456 | 1.000 | 0.736 | +0.264 | 0.706 | +0.294 |
| BTK and MAPK1 inputs to STAT5A removed | TP53 on | 0.373 | 0.571 | 0.487 | +0.084 | 0.483 | +0.088 |
| BTK and MAPK1 inputs to STAT5A removed | CDKN2A on | 0.482 | 0.764 | 0.669 | +0.095 | 0.654 | +0.110 |
| BTK and MAPK1 inputs to STAT5A removed | MYC off | 0.435 | 0.648 | 0.521 | +0.127 | 0.521 | +0.127 |
| BTK and MAPK1 inputs to STAT5A removed | CCND1 off | 0.541 | 0.578 | 0.503 | +0.075 | 0.521 | +0.057 |
| BTK and MAPK1 inputs to STAT5A removed | STAT5A off | 0.499 | 1.000 | 1.000 | +0.000 | 1.000 | +0.000 |

**Table 19:** A harder audit of the guarantee. Fifteen AML gene sets with the highest screening scores (many near-ties), *ε* = 0.02, *δ* = 0.05, true scores from 60,000 simulations each.

| Runs | Sets | Sets within 0.02 of best | Certified | ε-optimal | Interval violations | Mean simulations |
|---|---|---|---|---|---|---|
| 15 | 15 | 2 | 15 | 15 | 0 | 77,447 |

We also varied the per-node update noise of the Calzone network (sets of up to two nodes, *K* = 351; noise 0.005, 0.02 and 0.05). Certification and correctness held at every level (9/9 certified, 9/9 *ε*-optimal), with reductions of 6.8, 10.3 and 8.6*×* (Fig. 9a; Supplementary Table 15). At the highest noise level the achievable gain is smaller: the best set scored 0.724 against 0.577 untreated.

**Table 20:** All method-comparison runs. Decision rounds are the number of times the stopping rule is evaluated; for certified search each round is one parallel batch over all active candidates, for LUCB and UGapE each round is sequential.

| Family | ε | Method | Runs | Certified | ε-optimal | Interval violations | Mean simulations | SD | Mean decision rounds |
|---|---|---|---|---|---|---|---|---|---|
| AML druggable sets (K=92) | 0.01 | Certified search | 100 | 100 | 100 | 0 | 79,762 | 5,187 | 40 |
| AML druggable sets (K=92) | 0.01 | LUCB | 100 | 100 | 100 | 0 | 48,072 | 3,491 | 75 |
| AML druggable sets (K=92) | 0.01 | UGapE | 100 | 100 | 100 | 0 | 47,358 | 3,757 | 146 |
| AML druggable sets (K=92) | 0.01 | Doubling checkpoints | 100 | 100 | 100 | 0 | 92,732 | 11,346 | 6 |
| AML druggable sets (K=92) | 0.01 | Hoeffding radius | 100 | 100 | 100 | 0 | 156,330 | 10,298 | 134 |
| AML druggable sets (K=92) | 0.01 | No elimination | 100 | 100 | 100 | 0 | 747,408 | 94,865 | 41 |
| AML druggable sets (K=92) | 0.02 | Certified search | 100 | 100 | 100 | 0 | 71,564 | 4,649 | 26 |
| AML druggable sets (K=92) | 0.02 | LUCB | 100 | 100 | 100 | 0 | 39,224 | 1,935 | 53 |
| AML druggable sets (K=92) | 0.02 | UGapE | 100 | 100 | 100 | 0 | 38,492 | 2,251 | 101 |
| AML druggable sets (K=92) | 0.02 | Doubling checkpoints | 100 | 100 | 100 | 0 | 82,658 | 4,753 | 6 |
| AML druggable sets (K=92) | 0.02 | Hoeffding radius | 100 | 100 | 100 | 0 | 122,654 | 8,221 | 78 |
| AML druggable sets (K=92) | 0.02 | No elimination | 100 | 100 | 100 | 0 | 459,448 | 66,494 | 25 |
| AML druggable sets (K=92) | 0.05 | Certified search | 100 | 100 | 100 | 0 | 62,586 | 3,496 | 12 |
| AML druggable sets (K=92) | 0.05 | LUCB | 100 | 100 | 100 | 0 | 28,668 | 832 | 27 |
| AML druggable sets (K=92) | 0.05 | UGapE | 100 | 100 | 100 | 0 | 28,158 | 1,023 | 50 |
| AML druggable sets (K=92) | 0.05 | Doubling checkpoints | 100 | 100 | 100 | 0 | 74,178 | 4,993 | 5 |
| AML druggable sets (K=92) | 0.05 | Hoeffding radius | 100 | 100 | 100 | 0 | 91,560 | 6,614 | 26 |
| AML druggable sets (K=92) | 0.05 | No elimination | 100 | 100 | 100 | 0 | 210,312 | 26,831 | 11 |
| AML 3-gene sets (K=833) | 0.02 | Certified search | 50 | 50 | 50 | 0 | 597,664 | 28,745 | 242 |
| AML 3-gene sets (K=833) | 0.02 | LUCB | 50 | 50 | 50 | 0 | 274,948 | 7,167 | 1401 |
| AML 3-gene sets (K=833) | 0.02 | UGapE | 50 | 50 | 50 | 0 | 264,356 | 5,075 | 1812 |
| AML 3-gene sets (K=833) | 0.02 | Doubling checkpoints | 50 | 50 | 50 | 0 | 748,908 | 31,428 | 9 |
| AML 3-gene sets (K=833) | 0.02 | Hoeffding radius | 50 | 0 | 50 | 0 | 795,296 | 38,158 | 400 |
| AML 3-gene sets (K=833) | 0.02 | No elimination | 50 | 50 | 50 | 0 | 20,356,854 | 2,064,634 | 244 |
| AML 3-gene sets (K=833) | 0.05 | Certified search | 50 | 50 | 50 | 0 | 546,998 | 25,715 | 50 |
| AML 3-gene sets (K=833) | 0.05 | LUCB | 50 | 50 | 50 | 0 | 226,748 | 6,125 | 1003 |
| AML 3-gene sets (K=833) | 0.05 | UGapE | 50 | 50 | 50 | 0 | 205,846 | 2,434 | 1226 |
| AML 3-gene sets (K=833) | 0.05 | Doubling checkpoints | 50 | 50 | 50 | 0 | 691,568 | 33,473 | 7 |
| AML 3-gene sets (K=833) | 0.05 | Hoeffding radius | 50 | 50 | 50 | 0 | 710,884 | 32,687 | 152 |
| AML 3-gene sets (K=833) | 0.05 | No elimination | 50 | 50 | 50 | 0 | 4,184,992 | 346,353 | 50 |
| AML top-15 gene sets | 0.01 | Certified search | 100 | 1 | 100 | 0 | 123,269 | 4,110 | 400 |
| AML top-15 gene sets | 0.01 | LUCB | 100 | 0 | 100 | 0 | 91,882 | 979 | 478 |
| AML top-15 gene sets | 0.01 | UGapE | 100 | 0 | 100 | 0 | 91,944 | 779 | 905 |
| AML top-15 gene sets | 0.01 | Doubling checkpoints | 100 | 2 | 100 | 0 | 97,488 | 5,222 | 9 |
| AML top-15 gene sets | 0.01 | Hoeffding radius | 100 | 0 | 100 | 0 | 195,957 | 6,107 | 400 |
| AML top-15 gene sets | 0.01 | No elimination | 100 | 0 | 100 | 0 | 600,000 | 0 | 400 |
| AML top-15 gene sets | 0.02 | Certified search | 100 | 100 | 100 | 0 | 81,126 | 6,799 | 191 |
| AML top-15 gene sets | 0.02 | LUCB | 100 | 100 | 100 | 0 | 50,030 | 4,343 | 244 |
| AML top-15 gene sets | 0.02 | UGapE | 100 | 100 | 100 | 0 | 49,189 | 4,583 | 478 |
| AML top-15 gene sets | 0.02 | Doubling checkpoints | 100 | 100 | 100 | 0 | 78,128 | 13,416 | 8 |
| AML top-15 gene sets | 0.02 | Hoeffding radius | 100 | 0 | 100 | 0 | 196,007 | 5,761 | 400 |
| AML top-15 gene sets | 0.02 | No elimination | 100 | 100 | 100 | 0 | 289,050 | 36,546 | 193 |
| Calzone cell fate (K=2,951) | 0.02 | Certified search | 30 | 30 | 30 | 0 | 3,321,077 | 146,182 | 347 |
| Calzone cell fate (K=2,951) | 0.02 | LUCB | 30 | 30 | 30 | 0 | 1,146,223 | 23,309 | 7881 |
| Calzone cell fate (K=2,951) | 0.02 | UGapE | 30 | 30 | 30 | 0 | 1,151,193 | 15,046 | 8562 |
| Calzone cell fate (K=2,951) | 0.02 | Doubling checkpoints | 30 | 20 | 30 | 0 | 3,580,467 | 164,483 | 9 |
| Calzone cell fate (K=2,951) | 0.02 | Hoeffding radius | 30 | 0 | 30 | 0 | 4,517,497 | 140,237 | 400 |
| Calzone cell fate (K=2,951) | 0.02 | No elimination | 30 | 30 | 30 | 0 | 97,943,690 | 13,484,120 | 332 |
| Calzone cell fate (K=2,951) | 0.05 | Certified search | 30 | 30 | 30 | 0 | 2,560,553 | 101,733 | 65 |
| Calzone cell fate (K=2,951) | 0.05 | LUCB | 30 | 30 | 30 | 0 | 818,310 | 15,130 | 4697 |
| Calzone cell fate (K=2,951) | 0.05 | UGapE | 30 | 30 | 30 | 0 | 897,380 | 18,476 | 6024 |
| Calzone cell fate (K=2,951) | 0.05 | Doubling checkpoints | 30 | 30 | 30 | 0 | 2,956,287 | 108,381 | 7 |
| Calzone cell fate (K=2,951) | 0.05 | Hoeffding radius | 30 | 30 | 30 | 0 | 3,692,580 | 150,649 | 163 |
| Calzone cell fate (K=2,951) | 0.05 | No elimination | 30 | 30 | 30 | 0 | 18,866,727 | 1,848,485 | 64 |
| Grieco MAPK (K=1,275) | 0.02 | Certified search | 30 | 30 | 30 | 0 | 1,388,680 | 49,778 | 227 |
| Grieco MAPK (K=1,275) | 0.02 | LUCB | 30 | 30 | 30 | 0 | 583,917 | 22,198 | 3855 |
| Grieco MAPK (K=1,275) | 0.02 | UGapE | 30 | 30 | 30 | 0 | 589,157 | 14,812 | 4618 |
| Grieco MAPK (K=1,275) | 0.02 | Doubling checkpoints | 30 | 30 | 30 | 0 | 1,661,247 | 73,957 | 9 |
| Grieco MAPK (K=1,275) | 0.02 | Hoeffding radius | 30 | 0 | 30 | 0 | 2,260,997 | 69,504 | 400 |
| Grieco MAPK (K=1,275) | 0.02 | No elimination | 30 | 30 | 30 | 0 | 28,730,000 | 3,768,205 | 225 |
| Grieco MAPK (K=1,275) | 0.05 | Certified search | 30 | 30 | 30 | 0 | 1,209,453 | 51,364 | 50 |
| Grieco MAPK (K=1,275) | 0.05 | LUCB | 30 | 30 | 30 | 0 | 422,397 | 18,836 | 2394 |
| Grieco MAPK (K=1,275) | 0.05 | UGapE | 30 | 30 | 30 | 0 | 434,943 | 7,770 | 3075 |
| Grieco MAPK (K=1,275) | 0.05 | Doubling checkpoints | 30 | 30 | 30 | 0 | 1,469,453 | 63,375 | 7 |
| Grieco MAPK (K=1,275) | 0.05 | Hoeffding radius | 30 | 30 | 30 | 0 | 1,959,487 | 64,507 | 155 |
| Grieco MAPK (K=1,275) | 0.05 | No elimination | 30 | 30 | 30 | 0 | 6,515,250 | 455,851 | 51 |
| Remy tumorigenesis (K=5,488) | 0.02 | Certified search | 20 | 20 | 20 | 0 | 13,161,835 | 500,574 | 86 |
| Remy tumorigenesis (K=5,488) | 0.02 | LUCB | 20 | 20 | 20 | 0 | 4,193,530 | 38,175 | 35837 |
| Remy tumorigenesis (K=5,488) | 0.02 | UGapE | 20 | 20 | 20 | 0 | 5,145,780 | 103,748 | 45971 |
| Remy tumorigenesis (K=5,488) | 0.02 | Doubling checkpoints | 20 | 20 | 20 | 0 | 16,933,180 | 2,029,410 | 8 |
| Remy tumorigenesis (K=5,488) | 0.02 | Hoeffding radius | 20 | 0 | 20 | 0 | 51,781,700 | 233,607 | 400 |
| Remy tumorigenesis (K=5,488) | 0.02 | No elimination | 20 | 20 | 20 | 0 | 46,565,680 | 4,065,236 | 85 |
| Remy tumorigenesis (K=5,488) | 0.05 | Certified search | 20 | 20 | 20 | 0 | 7,840,070 | 277,128 | 30 |
| Remy tumorigenesis (K=5,488) | 0.05 | LUCB | 20 | 20 | 20 | 0 | 2,564,075 | 37,615 | 19346 |
| Remy tumorigenesis (K=5,488) | 0.05 | UGapE | 20 | 20 | 20 | 0 | 3,462,525 | 43,972 | 29138 |
| Remy tumorigenesis (K=5,488) | 0.05 | Doubling checkpoints | 20 | 20 | 20 | 0 | 9,124,420 | 97,217 | 6 |
| Remy tumorigenesis (K=5,488) | 0.05 | Hoeffding radius | 20 | 20 | 20 | 0 | 27,355,555 | 828,902 | 154 |
| Remy tumorigenesis (K=5,488) | 0.05 | No elimination | 20 | 20 | 20 | 0 | 16,271,920 | 1,198,068 | 30 |

**Table 21:** Effect of the batch size of certified search. Replayed pools; *ε* = 0.01 (*K* = 92, 60 runs per size) and *ε* = 0.02 (*K* = 833, 30 runs per size).

| Family | Batch size *m* | Mean simulations | Certified | Interval violations |
|---|---|---|---|---|
| AML 3-gene sets (K=833) | 50 | 639,532 | 30/30 | 0 |
| AML 3-gene sets (K=833) | 100 | 598,490 | 30/30 | 0 |
| AML 3-gene sets (K=833) | 200 | 585,653 | 30/30 | 0 |
| AML 3-gene sets (K=833) | 400 | 597,707 | 30/30 | 0 |
| AML 3-gene sets (K=833) | 1000 | 911,100 | 30/30 | 0 |
| AML druggable sets (K=92) | 50 | 90,333 | 60/60 | 0 |
| AML druggable sets (K=92) | 100 | 85,848 | 60/60 | 0 |
| AML druggable sets (K=92) | 200 | 79,430 | 60/60 | 0 |
| AML druggable sets (K=92) | 400 | 77,060 | 60/60 | 0 |
| AML druggable sets (K=92) | 1000 | 112,767 | 60/60 | 0 |

**Table 22:** Controlled test of family size and near-ties. Families of *K* candidate sets are drawn from the 833 three-gene sets; the *j* highest-scoring sets are always included and the rest are random (five sub-families per row, six runs each, *ε* = 0.02). *H*_*ε*_ = ∑_*a*_ max(Δ_*a*_, *ε*)^*−*2^.

| *K* | Best sets always included *j* | Mean gap complexity *H*<sub>*ε*</sub> | Mean simulations | Uniform, same guarantee | Reduction | Certified |
|---|---|---|---|---|---|---|
| 50 | 5 | 8,218 | 87,517 | 475,100 | 5.4× | 100% |
| 100 | 1 | 5,801 | 63,010 | 1,036,800 | 16.5× | 100% |
| 100 | 3 | 8,986 | 113,030 | 1,036,800 | 9.2× | 100% |
| 100 | 5 | 9,439 | 118,000 | 1,036,800 | 8.8× | 100% |
| 100 | 10 | 10,266 | 127,023 | 1,036,800 | 8.2× | 100% |
| 100 | 20 | 11,268 | 141,153 | 1,036,800 | 7.3× | 100% |
| 200 | 5 | 11,793 | 178,717 | 2,246,800 | 12.6× | 100% |
| 400 | 5 | 17,062 | 309,443 | 4,840,400 | 15.6× | 100% |
| 833 | 5 | 27,817 | 602,530 | 10,843,994 | 18.0× | 100% |

**Table 23:** Elapsed time with parallel workers. Mean elapsed time in units of one batch-of-*m*-simulations per worker, for *W* = 16 workers (columns 3–6) and *W* = 128 workers (columns 7–10). A round of certified search runs its active candidates in parallel batches (⌈*A/W* ⌉ batch times for *A* active candidates); each LUCB or UGapE round is one batch time, because the next choice depends on the previous result. Dispatch latency is not charged, which favours the sequential methods. Serial time (*W* = 1) equals the total simulations of Table 2. Same runs as Table 2.

| Family | ε | Certified | LUCB | UGapE | Doubling | Certified | LUCB | UGapE | Doubling |
|---|---|---|---|---|---|---|---|---|---|
| AML druggable sets (K=92) | 0.01 | 10,766 | 16,036 | 30,158 | 12,452 | 7,978 | 15,036 | 29,158 | 9,280 |
| AML druggable sets (K=92) | 0.02 | 8,032 | 11,612 | 21,292 | 9,196 | 5,256 | 10,612 | 20,292 | 6,144 |
| AML druggable sets (K=92) | 0.05 | 5,094 | 6,334 | 10,958 | 6,200 | 2,314 | 5,334 | 9,958 | 2,992 |
| AML 3-gene sets (K=833) | 0.02 | 55,770 | 145,260 | 186,356 | 65,836 | 27,538 | 140,660 | 181,756 | 29,128 |
| AML 3-gene sets (K=833) | 0.05 | 36,672 | 105,516 | 127,846 | 46,668 | 8,354 | 100,916 | 123,246 | 10,616 |
| AML top-15 gene sets | 0.01 | 39,991 | 47,767 | 90,544 | 25,600 | 39,991 | 47,767 | 90,544 | 25,600 |
| AML top-15 gene sets | 0.02 | 19,131 | 24,365 | 47,789 | 16,128 | 19,131 | 24,365 | 47,789 | 16,128 |
| Calzone cell fate (K=2,951) | 0.02 | 225,133 | 806,467 | 874,593 | 234,640 | 51,330 | 790,367 | 858,493 | 44,653 |
| Calzone cell fate (K=2,951) | 0.05 | 163,097 | 488,073 | 620,780 | 187,873 | 23,060 | 471,973 | 604,680 | 25,387 |
| Grieco MAPK (K=1,275) | 0.02 | 95,630 | 393,447 | 469,657 | 113,773 | 29,950 | 386,447 | 462,657 | 34,773 |
| Grieco MAPK (K=1,275) | 0.05 | 78,017 | 247,337 | 315,443 | 94,673 | 12,310 | 240,337 | 308,443 | 15,947 |
| Remy tumorigenesis (K=5,488) | 0.02 | 826,450 | 3,617,880 | 4,631,280 | 1,063,740 | 106,800 | 3,587,880 | 4,601,280 | 136,260 |
| Remy tumorigenesis (K=5,488) | 0.05 | 491,330 | 1,968,840 | 2,948,025 | 571,560 | 62,625 | 1,938,840 | 2,918,025 | 72,740 |

**Table 24:** Out-of-sample agreement of the calibrated network with measured single-drug response (public data only). Spearman correlation between the network output and measured BeatAML AUC (FPMTB: sensitivity score, sign reversed). Held-out drug: five folds, each fitted without one drug (test *n* = 8 genotypes). Held-out genotype: eight folds, each fitted without one mutation profile (*n* = 5 drugs). Brackets: bootstrap 95% interval. Trivial baseline: mean AUC of the other drugs at the same genotype (held-out drug) or of the other genotypes for the same drug (held-out genotype). Persistence: the measured training-cohort values themselves, which a perfect fit to the training cohort would reproduce (FPMTB: drug-level values only). No combination data are involved.

| Analysis | Test set | *n* | Model Spearman *ρ* | Permutation *p* | Baseline *ρ* | Baseline |
|---|---|---|---|---|---|---|
| Held-out drug | Venetoclax | 8 | 0.67 | 0.083 | 0.40 | trivial |
| Held-out drug | Ibrutinib | 8 | 0.00 | 1.000 | 0.67 | trivial |
| Held-out drug | Entospletinib | 8 | -0.55 | 0.171 | 0.69 | trivial |
| Held-out drug | Selumetinib | 8 | -0.33 | 0.428 | -0.43 | trivial |
| Held-out drug | Midostaurin | 8 | 0.37 | 0.360 | 0.55 | trivial |
| Held-out drug | pooled | 40 | 0.28 [-0.05, 0.59] | 0.079 | 0.11 | trivial |
| Held-out genotype | pooled | 40 | 0.28 [-0.04, 0.57] | 0.078 | 0.37 | trivial |
| Cross-cohort | Waves 3+4 fit → Waves 1+2 data | 40 | 0.65 [0.43, 0.80] | ¡0.001 | 0.77 | persistence |
| Cross-cohort | Waves 1+2 fit → Waves 3+4 data | 40 | 0.69 [0.46, 0.84] | ¡0.001 | 0.77 | persistence |
| Cross-cohort | Waves 3+4 fit → FPMTB cohort (sign reversed) | 22 | 0.60 [0.21, 0.85] | 0.004 | 0.61 | persistence |
| Cross-cohort | Waves 1+2 fit → FPMTB cohort (sign reversed) | 22 | 0.56 [0.18, 0.77] | 0.007 | 0.61 | persistence |

**Table 25:** All decoy statistics, with raw and multiplicity-adjusted *p*-values. Four statistics were computed on the same decoy maps; none was designated primary in advance, and all four are shown. Raw *p* = (1 + #*{*decoys ≥ real*}*)*/*(1 + #decoys) using the mean of the replicate evaluations of the real map. Holm adjustment is over the four statistics. The last column gives the raw *p* obtained from each single replicate of the real map, which shows how much a single evaluation of the real map would move the result. The first statistic is the most conservative because every decoy map also benefits from selecting the largest excess among its own sets.

| Statistic | Real map | Decoy median | Decoy 95th pct | Decoys ≥ real | Raw *p* | Holm *p* | Raw *p* range over replicates |
|---|---|---|---|---|---|---|---|
| Largest excess over additivity among all 92 sets (selection-aware) | 0.152 | 0.066 | 0.148 | 42/1000 | 0.043 | 0.070 | 0.014–0.158 |
| Excess over additivity of the best-scoring set | 0.152 | 0.032 | 0.118 | 5/1000 | 0.006 | 0.020 | 0.002–0.050 |
| Excess over Bliss independence of the best-scoring set | 0.161 | 0.037 | 0.131 | 4/1000 | 0.005 | 0.020 | 0.002–0.051 |
| Score of the best-scoring set | 0.712 | 0.554 | 0.697 | 34/1000 | 0.035 | 0.070 | 0.012–0.057 |

**Table 26:** Single-gene loss of function. Each row holds one gene off throughout (also when untreated) and repeats the 92-set search with 1,500 simulations per set. The triplet is ibrutinib + midostaurin + ERK-inhibitor class (ERKi); FLT3i: gilteritinib-class FLT3 inhibitor. Reward genes and the mutation nodes were not removed. Ranks of near-tied sets can change between runs because of simulation noise.

| Gene held off | Best set | Triplet rank | Triplet score | Triplet excess (additive) |
|---|---|---|---|---|
| none (reference) | Ibrutinib + Midostaurin + ERKi | 1 | 0.707 | +0.131 |
| AKT1 | Ibrutinib + Midostaurin + ERKi | 1 | 0.707 | +0.100 |
| BCL2 | Ibrutinib + Midostaurin + ERKi | 1 | 0.707 | +0.131 |
| BTK | Midostaurin + ERKi | 6 | 0.707 | +0.090 |
| CEBPA | Ibrutinib + FLT3i + ERKi | 2 | 0.617 | +0.094 |
| CTNNB1 | Ibrutinib + Entospletinib + ERKi | 3 | 0.726 | +0.117 |
| ETV6 | Ibrutinib + Midostaurin + ERKi | 1 | 0.707 | +0.131 |
| FBXW7 | Ibrutinib + FLT3i + ERKi | 3 | 0.609 | +0.088 |
| FLT3 | Ibrutinib + Midostaurin + ERKi | 1 | 0.707 | +0.048 |
| GSK3B | Ibrutinib + FLT3i + ERKi | 3 | 0.582 | +0.066 |
| HOXA9 | Ibrutinib + Midostaurin + ERKi | 1 | 0.707 | +0.131 |
| KIT | Ibrutinib + Entospletinib + ERKi | 3 | 0.707 | +0.118 |
| MAP2K1 | Ibrutinib + FLT3i + ERKi | 3 | 0.589 | +0.068 |
| MAP2K2 | Ibrutinib + Midostaurin + ERKi | 1 | 0.707 | +0.118 |
| MAPK1 | Ibrutinib + Midostaurin + FLT3i | 5 | 0.756 | +0.045 |
| MEIS1 | Ibrutinib + Midostaurin + ERKi | 1 | 0.707 | +0.131 |
| SOX4 | Ibrutinib + Midostaurin + ERKi | 1 | 0.710 | +0.131 |
| SYK | Ibrutinib + Midostaurin + ERKi | 1 | 0.707 | +0.036 |

### Supplementary Note 7: reproducing the results

All analyses are in one Python file (`aml_certified search.py`). It runs on a local machine or in Google Colab and writes every table and figure to a results folder; seeds are fixed. The file is organised in stages that can be run separately, and the AML clamping policy can be switched with `--aml-policy` (`strict`, the default, or `legacy`). Stages that take longer than a few minutes must be allowed to finish in one session; if a run is interrupted, re-run the remaining stages. The numbers in the text are generated from the result tables by a script, which ties the text to the data. During the revision we replaced a process-dependent random seed by a fixed one; the reported values are from runs with the fixed seeds. The reported results were produced over several sessions because long runs were interrupted. The out-of-sample check (Supplementary Note 8) uses only files shipped with KGBN, and the decoy and loss-of-function tests (Supplementary Note 9) use only the network; neither depends on the clamping policy. Parts D.1 (audit) and D.2 (sensitivity), the Calzone noise analysis, the patient-level analysis, and the Calzone, Grieco and Remy benchmark tables come from earlier complete runs of the same code (they do not depend on the AML clamping policy); the AML results, the extended benchmark for the AML network (eight repetitions) and all new analyses were produced after the policy correction. We have not repeated the complete pipeline in one uninterrupted run, and we make no claim of bit-for-bit reproducibility across library versions.

### Supplementary Note 8: out-of-sample check of the calibrated network

This note reports a retrospective check on public data that were already collected; no new data were generated. It tests whether the fitted network predicts *single-drug* response in data it was not fitted to. It cannot test combinations, because no combination data were used. Table 24 and Fig. S3 give the results and the main text interprets them. Points to bear in mind: the samples are small, the cross-cohort agreement is dominated by differences between drugs, the cohorts overlap in the measured quantity, and the toolkit’s fitting procedure was not designed to answer our question. A correlation with held-out single-drug data would in any case be necessary but not sufficient for trusting a combination prediction.

**Figure S3:**
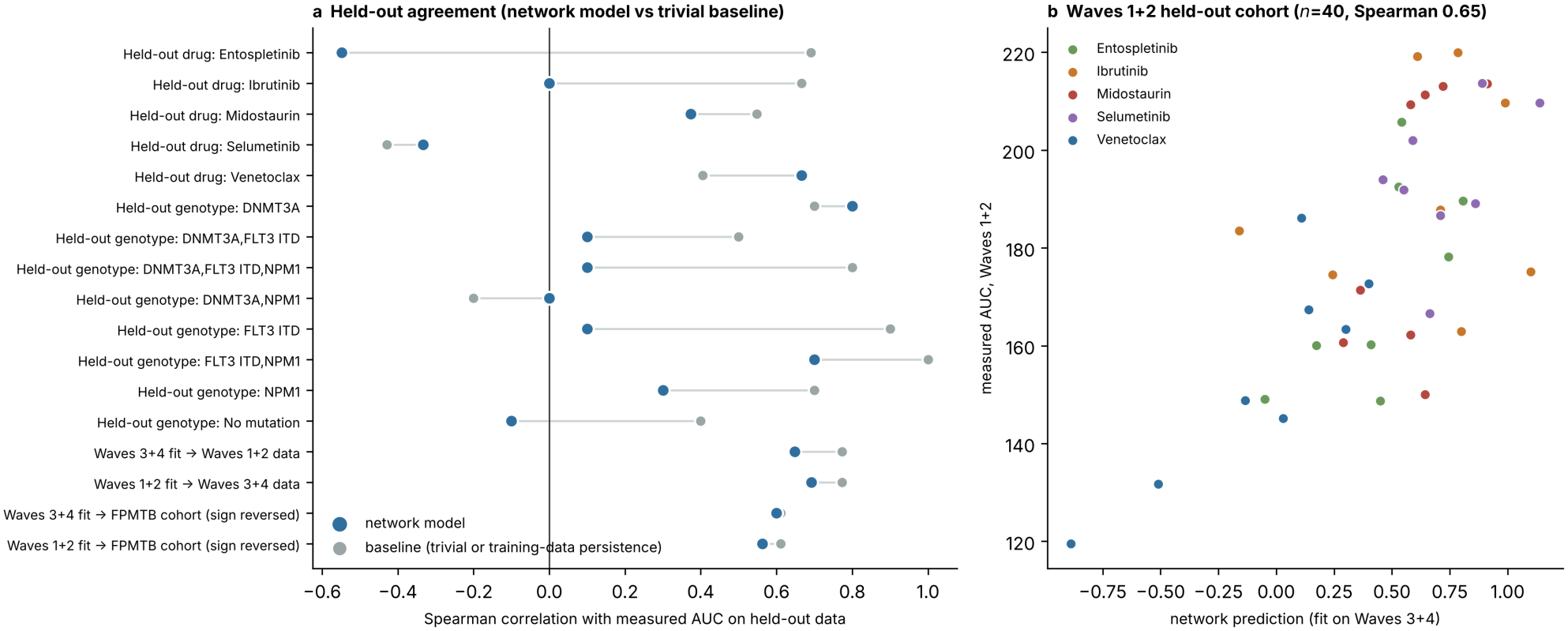
Held-out agreement of the calibrated network. **a**, Spearman correlation with measured AUC for each held-out drug, each held-out mutation profile and each cross-cohort evaluation, for the network (blue) and the baseline (grey; trivial baseline for held-out folds, measured training-cohort values for cross-cohort rows; FPMTB sign reversed). **b**, Network prediction against measured AUC in Waves 1+2 for the Waves 3+4 fit.

### Supplementary Note 9: decoy drug-target maps and single-gene loss of function

Both tests are computational and use only the network. The decoy test asks whether a result as strong as the real one needs the real drug-to-gene map. It is a test of the model, not of biology: a small *p*-value means that random maps rarely give a best set this good and this super-additive, not that the real map is right. The decoy median is positive (0.032) because the search selects the best of 92 sets, so some excess is expected by selection alone. Decoys with the highest scores exist (Fig. S4a), so the real map is not at the extreme. Table 25 gives all four statistics. The first is the most conservative and is the one we would regard as the primary comparison in hindsight, but it was chosen after seeing the others and we therefore report all four with the Holm adjustment. Because the raw *p*-values depend on the noise in the single estimate of the real map (last column), we used 30 replicate evaluations of the real map and 1000 decoys. For transparency: an earlier run with 300 decoys and 10 replicates gave raw *p* = 0.010 for the excess over additivity of the best-scoring set and 0.056 for the selection-aware statistic. We then increased the numbers to reduce Monte Carlo error before seeing the larger run’s results, and we report only the larger run, which gives different values for the same statistics (Table 25); the difference illustrates how unstable single *p*-values from this procedure are. The loss-of-function test removes one gene at a time and repeats the search with fewer simulations per set, so ranks of near-tied sets can change between runs.

**Figure S4:**
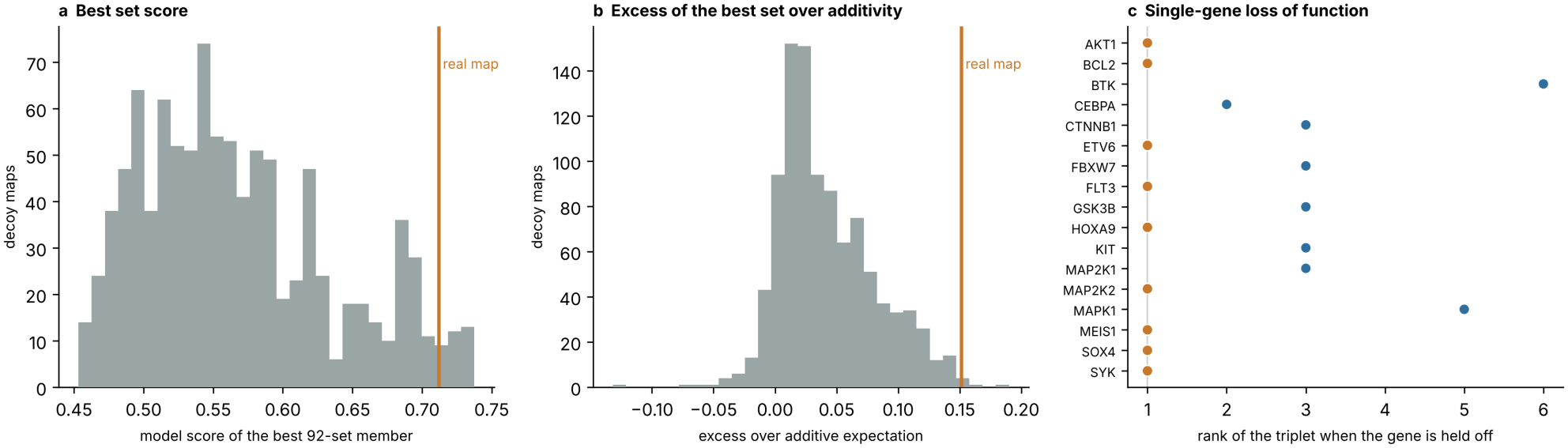
Decoy drug-target maps and single-gene loss of function. **a**, Score of the best set in the 92-set search under 1000 decoy maps (grey) and under the real map (orange line). **b**, Excess of the best set over additivity, same maps. **c**, Rank of the ibrutinib + midostaurin + ERK-inhibitor triplet when each gene is held off (orange: rank 1).

## Notes

https://doi.org/10.5281/zenodo.24450270

https://github.com/hklarner/pyboolnet

https://github.com/IlyaLab/KGBN

https://github.com/wasimrazaa135-ops/aml-certified-search

### Competing Interest Statement

The authors have declared no competing interest.

